# The derlin Dfm1 promotes retrotranslocation of folded protein domains from the endoplasmic reticulum

**DOI:** 10.1101/2020.06.02.131128

**Authors:** Daniel Fonseca, Pedro Carvalho

**Affiliations:** Sir William Dunn School of Pathology, University of Oxford, South Parks Road, Oxford OX1 3RE, UK

## Abstract

Endoplasmic reticulum (ER) proteins are degraded by proteasomes in the cytosol through ER-associated degradation (ERAD). This process involves retrotranslocation of substrates across the ER membrane, their ubiquitination and subsequent membrane extraction by the Cdc48/Npl4/Ufd1 ATPase complex prior delivery to proteasomes for degradation. Recently a mechanism for the retrotranslocation of misfolded substrates by the Hrd1 ubiquitin ligase complex was described. However, how substrates with folded luminal domains are retrotranslocated remains unknown. Here, we identify Dfm1 as an essential membrane component for the retrotranslocation of proteins with folded luminal domains. Both Dfm1 intramembrane rhomboid-like and the cytosolic Cdc48-binding domains are essential for substrate retrotranslocation. Substrate processing by Dfm1 and Cdc48 complex requires the ubiquitin shuttle factors Rad23/Dsk2 and the multi-ubiquitination enzyme Ufd2. Our findings suggest a pathway in which a series of ubiquitin modifying factors recruit Dfm1 to resolve a stalled retrotranslocation intermediate due to the presence of a folded luminal domain.

## INTRODUCTION

Proteins in the endoplasmic reticulum (ER) are degraded by ER-associated protein degradation (ERAD) (Christianson and Ye, 2014; Ruggiano et al., 2014). This process targets a broad range of substrates, including misfolded membrane and secretory proteins generated as byproducts of biosynthetic processes, as well as a set of folded ER proteins, mostly rate-limiting enzymes of lipid metabolism as part of homeostatic regulatory mechanisms. Thus, ERAD controls both protein and lipid homeostasis in the ER and has a central role in cellular physiology in health and disease (Bhattacharya and Qi, 2019; Needham et al., 2019).

ERAD is carried out by multiple, membrane integral ubiquitin ligase complexes, each with specificity for distinct substrate classes (Christianson and Ye, 2014; Ruggiano et al., 2014). In yeast, there are three ERAD ubiquitin ligase complexes -the Hrd1, Doa10 and Asi complexes- and despite their different substrate specificities, the general steps leading to substrate degradation are similar (Wu and Rapoport, 2018). Upon recognition and engagement with an ERAD ubiquitin ligase complex, substrates are transported across the ER membrane into the cytosol, a process called retrotranslocation. Once exposed on the cytosolic surface of the ER, substrates are polyubiquitinated and subsequently extracted from the ER membrane through the activity of the Cdc48 complex, which includes the hexameric Cdc48 ATPase as well as Npl4 and Ufd1 cofactors (Bays et al., 2001; Ye et al., 2001; Jarosch et al., 2002; Stein et al., 2014; Bodnar and Rapoport, 2017). A number of adaptor proteins, including the multi-ubiquitination enzyme Ufd2 and the shuttle factors Rad23 and Dsk2, finally hand the ubiquitinated substrate to the proteasome for degradation (Medicherla et al., 2004; Richly et al., 2005; Baek et al., 2011).

Despite this general scheme, the molecular mechanisms of the individual ERAD steps are less clear, in particular substrate retrotranslocation across the ER membrane. *In vivo* and *in vitro* studies suggest that for ER luminal substrates, the major retrotranslocation component is the Hrd1 ubiquitin ligase (Carvalho et al., 2010; Baldridge and Rapoport, 2016; Schoebel et al., 2017; Vasic et al., 2020; Wu et al., 2020). Hrd1-mediated retrotranslocation depends on its membrane domain, which recent cryo-EM studies revealed to display features reminiscent of other protein-conducting channels such as the Sec translocon that transports unfolded, newly synthesized proteins into the ER lumen (Schoebel et al., 2017; Wu et al., 2020). The yeast derlins Der1 and Dfm1, members of the rhomboid protein superfamily (Freeman, 2014), have also been implicated in retrotranslocation (Mehnert et al., 2014; Neal et al., 2018; Wu et al., 2020). Despite their similarity, Der1 and Dfm1 appeared to have distinct substrate specificity. While Der1 interacts with and is required for the degradation of misfolded ER luminal substrates (Knop et al., 1996; Carvalho et al., 2006; Denic et al., 2006), Dfm1 appears to target mostly misfolded membrane proteins (Stolz et al., 2010; Neal et al., 2018). The basis for these substrate preferences as well as the precise mechanisms by which ERAD components facilitate retrotranslocation remain obscure. The picture is even less clear for the retrotranslocation of ERAD substrates containing folded domains. It is well established that folded domains within the ER lumen slow down the degradation of ERAD substrates suggesting that their retrotranslocation is challenging (Tirosh et al., 2003; Bhamidipati et al., 2005; Carvalho et al., 2010; Shi et al., 2019). However, how such domains are retrotranslocated and whether the process requires additional components is unknown.

Most ERAD substrates characterized so far are degraded constitutively and exist in cells as an heterogeneous ensemble of intermediates. Constitutive degradation of substrates also precludes the uncoupling between protein folding and retrotranslocation. Here, we circumvented these problems by employing an inducible auxin-based degron system to study protein retrotranslocation. By uncoupling protein folding from subsequent steps, we searched for factors required for the retrotranslocation of luminal folded domains. We uncovered that the retrotranslocation of folded domains requires specifically the derlin Dfm1, while ERAD components necessary for the retrotranslocation of misfolded proteins, such as Hrd1 or Der1 are dispensable. Dfm1 function requires both its rhomboid-like membrane domain as well as its cytosolic interaction with the Cdc48/Npl4/Ufd1 ATPase complex. Finally, we show that a cascade of ubiquitin processing factors is necessary for retrotranslocation mediated by Dfm1 and the Cdc48 complex. These findings delineate a pathway for the resolution of halted retrotranslocation events due to the presence of folded luminal domains.

## RESULTS

### An auxin-induced assay to study membrane protein retrotranslocation

To study how folded luminal domains in membrane proteins are retrotranslocated for proteasomal degradation we generated a chimeric membrane protein with the tightly folded dihydrofolate reductase (DHFR) domain from *E. coli* in the ER lumen. Generation of DHFR fusion proteins is a well-established approach to study protein translocation across lipid bilayers (Eilers and Schatz, 1986; Arkowitz et al., 1993; Gambill et al., 1993; Tirosh et al., 2003; Bhamidipati et al., 2005; Shi et al., 2019). The chimeric membrane protein was engineered to adopt a type I topology in the ER membrane with the N-terminal signal sequence of the ER chaperone Kar2 followed by a triple HA tag, an opsin glycosylation site, the DHFR domain and the transmembrane segment of Ost1, an ER protein. The cytosolic domain of the chimera contained four tandem repeats of the 44-amino acid minimal degron (AID*) derived from the *Arabidopsis thaliana* IAA17 protein (Morawska and Ulrich, 2013) followed by nine repeats of the MYC epitope. The chimeric construct was termed inducible Retro-Clogger (iRC) for reasons explained below (Figure 1A).

**Figure 1.**
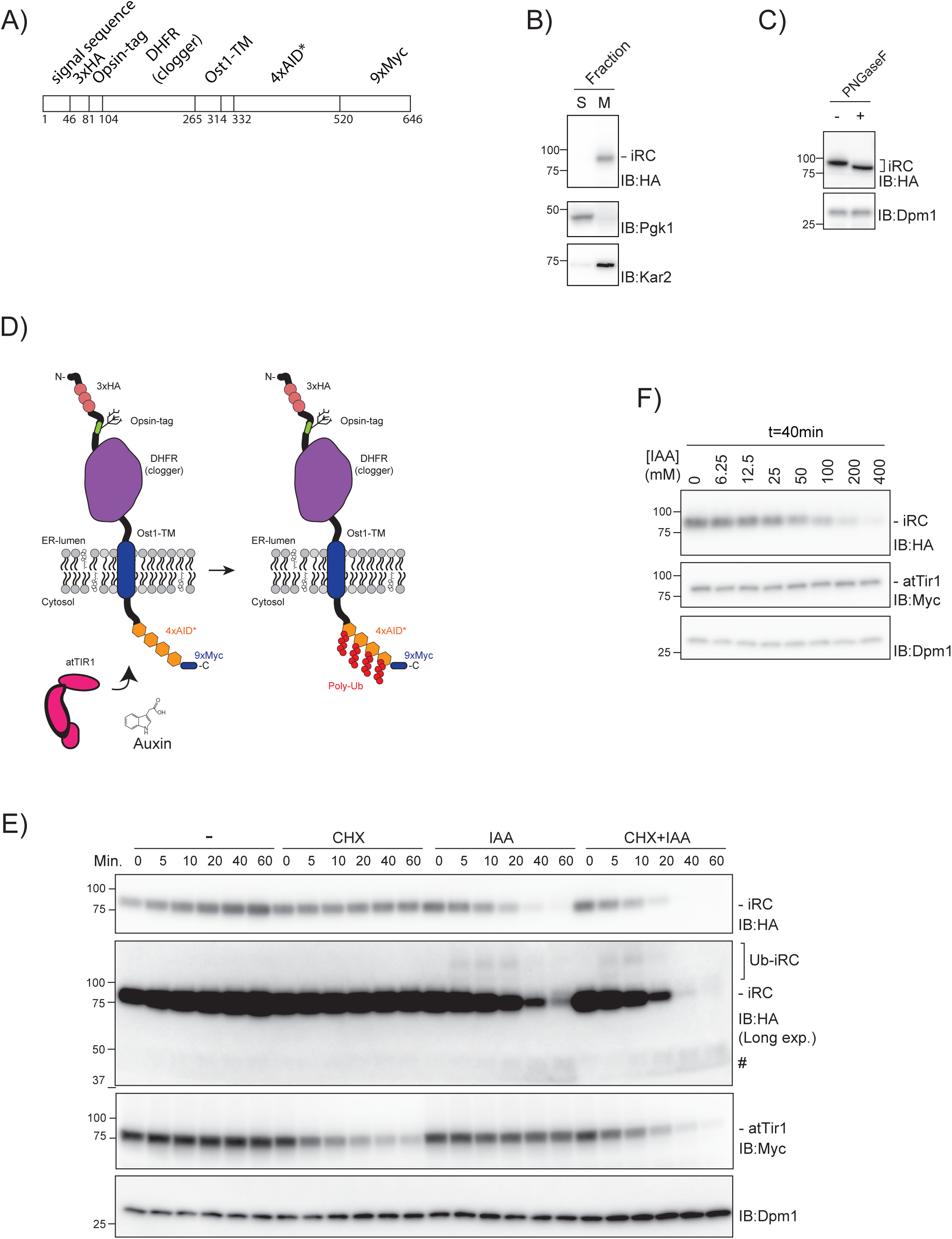
Inducible Retro-Clogger (iRC), an inducible substrate to study protein retrotranslocation. (A) Scheme of the inducible Retro-Clogger (iRC) substrate. The individual modules that compose the iRC are indicated. Numbers indicate the amino acid borders of each module. (B) iRC is membrane-associated. Crude membrane (M) and soluble (S) fractions from *WT* cells expressing iRC were prepared and analyzed by SDS-PAGE and western blotting using an α-HA antibody. The ER-associated protein Kar2 and the cytosolic phosphoglycerate kinase (Pgk1) proteins were used as controls and detected with α-Kar2 and Pgk1 antibodies, respectively. (C) The iRC adopts a type I topology with the N-terminus in the ER lumen. Whole-cell extracts of *WT* cells expressing iRC were treated with PNGase F and analyzed by SDS–PAGE and western blotting using an α-HA antibody. The non-glycosylated dolichol phosphate mannose synthase (Dpm1) was used as a loading control and detected with the α-Dpm1 antibody (D) Schematic representation of the topology of the iRC chimera and of its auxin-dependent ubiquitination. (E) Time-course analysis of iRC stability in cells expressing AtTir1 under the indicated conditions. Auxin (IAA) and cycloheximide (CHX) were added at time 0 and were used at 0.4mM and 125μg/ml, respectively. Note that addition of IAA leads to the appearance of faint high molecular weight smear consistent with iRC ubiquitination (Ub-iRC in Long exposure). There is also the appearance of a faint low molecular doublet in late timepoints after IAA addition (#). Whole-cell extracts were analyzed by SDS–PAGE and western blotting. iRC and AtTir1 were detected with α-HA and α-Myc antibodies, respectively. Dpm1 was used as a loading control and detected with the α-Dpm1 antibody. Timepoints are in minutes. (F) Dependence of auxin concentration in iRC degradation. Levels of iRC in *WT* cells expressing AtTir1 were analyzed after 40min incubation with the indicated concentration (in mM) of auxin (IAA). Samples prepared and analyzed as in (E).

Using fractionation experiments, we established that the iRC substrate was inserted in the ER membrane (Figure 1B). To confirm that iRC acquired the correct topology we examined whether iRC the opsin sequence was glycosylated. Treatment of yeast lysates with Peptide:N-glycosidase F (PNGaseF), which digest the N-glycans, resulted in faster iRC migration by SDS-PAGE (Figure 1C). Thus, iRC is targeted to the ER membrane with its N-terminus facing the ER lumen (Figure 1D).

ER ubiquitin ligases, such as Hrd1, are required for both ubiquitination and retrotranslocation of ERAD substrates (Carvalho et al., 2010; Stein et al., 2014; Baldridge and Rapoport, 2016; Vasic et al., 2020). To study substrate retrotranslocation bypassing the need of ubiquitination by the endogenous ER-resident ubiquitin ligases, we exploited the plant-based auxin-induced degradation (AID) (Nishimura et al., 2009; Morawska and Ulrich, 2013). The AID system relies on the expression of a plant-specific F-box protein, Tir1, to recognize AID* motifs in substrates only in the presence of auxin (indole-3-acetic acid, or IAA), triggering their polyubiquitination and degradation (Figure 1D).

In yeast cells expressing constitutively the *Arabidopsis thaliana* Tir1 (AtTir1) and in the absence of auxin, iRC is a long-lived protein. Even upon inhibition of protein synthesis with cycloheximide, there is no appreciable iRC turnover over a period of 60 minutes (Figure 1E). This was not due to ineffective translation inhibition as Tir1 protein levels decrease over time, indicating that it is a short-lived protein. Upon auxin addition, the iRC was rapidly degraded irrespective of the presence of cycloheximide (without cycloheximide t_1/2_=15min; with cycloheximide t_1/2_=10min). Importantly, iRC degradation was selective even at higher auxin concentrations (Figures 1E and F). Thus, auxin-induced ubiquitination of iRC is sufficient to trigger its retrotranslocation and degradation in an acute and synchronous manner.

Long exposures of the blots with the iRC chases upon auxin addition revealed two interesting features (Figure 1E). First, auxin addition resulted in high molecular weight smears suggestive of iRC ubiquitination. Second, while most iRC was degraded, auxin addition resulted in the appearance of small amounts of ∼40kDa fragments. These fragments did not contain the MYC epitope present at the C-terminal region of full-length iRC (see below). Thus, based on the size, they likely correspond to the tightly folded DHFR, suggesting that retrotranslocation of this domain is rate-limiting.

### iRC degradation does not require ERAD ubiquitin ligases

Increasing evidence implicates the ERAD ubiquitin ligases in substrate retrotranslocation (Carvalho et al., 2010; Stein et al., 2014; Baldridge and Rapoport, 2016; Natarajan et al., 2020; Wu et al., 2020). Therefore, we tested whether they were necessary for iRC retrotranslocation and degradation. Individual deletion of Asi1, Hrd1 and Doa10 did not prevent auxin-induced degradation of iRC (Figure 2A). Moreover, the degradation of iRC was unaffected by simultaneous deletion of the three ubiquitin ligases or the ubiquitin-conjugating enzyme Ubc7 involved in all ERAD branches (Figure 2A). As in *wild-type* (*WT*) cells, induction of iRC degradation in ERAD ubiquitin ligase mutants resulted in minute accumulation of the ∼40kDa fragment. These results indicate that auxin-induced degradation of iRC substrate is independent of the canonical ERAD ubiquitin ligases and does not require their retrotranslocation activity.

**Figure 2.**
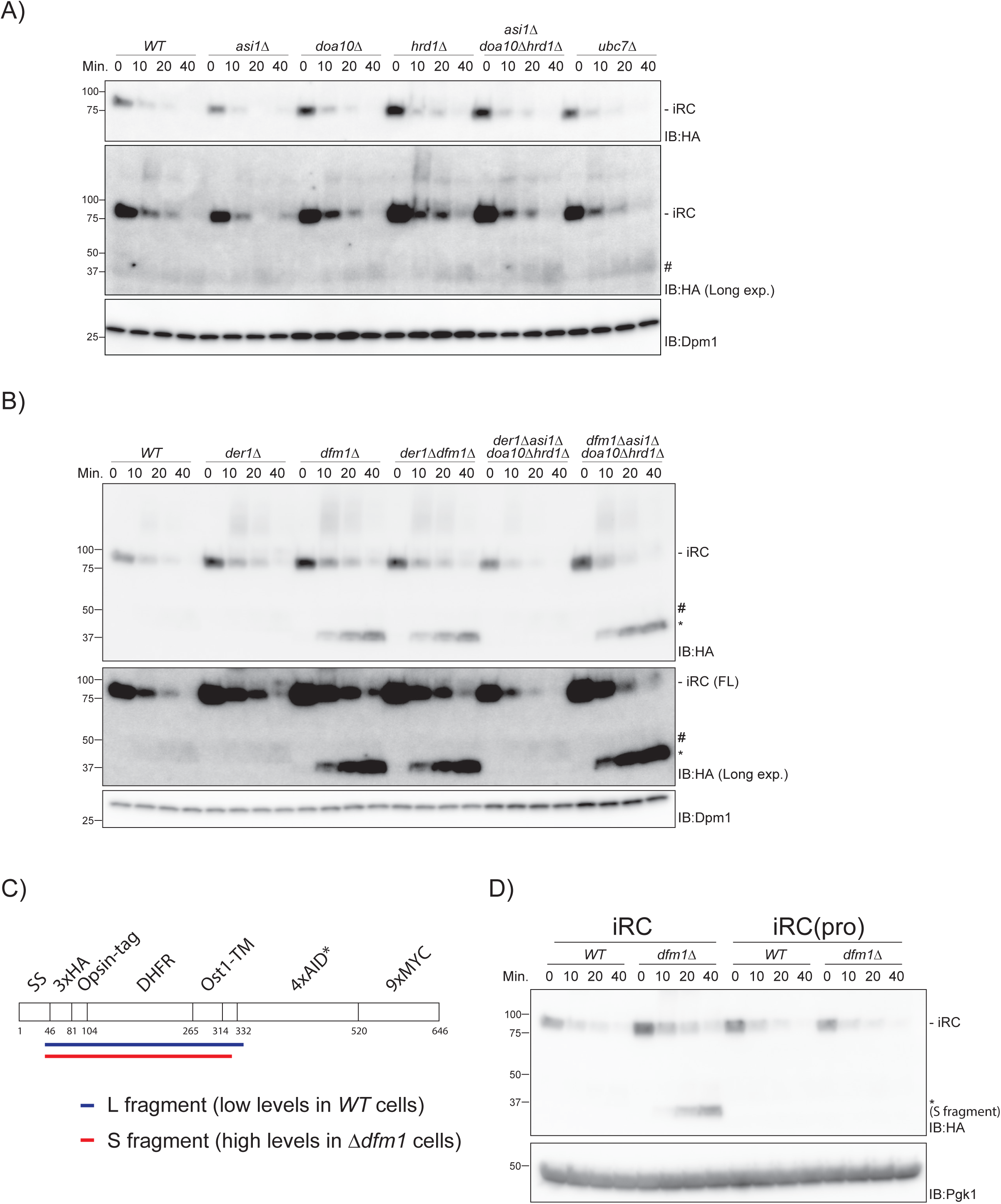
Complete iRC degradation requires Dfm1 but not other ERAD membrane components. (A) The degradation of iRC analyzed in cells lacking the indicated ERAD components, as in Figure 1 (E). (B) The iRC degradation was analyzed as in (A) in *WT* cells or cells with the indicated deletions. Note that iRC degradation in cells lacking Dfm1 lead to accumulation of a fragment of ∼37kDa (*). This fragment is present in substantially higher amounts and is slightly smaller than the one observed cells expressing Dfm1 (#). (C) Schematic representation of the iRC-derived L (long, in blue) and S (short, in red) fragments present in WT and *dfm1Δ* cells, respectively, as detected by mass spectrometry. Detergent extracts prepared from *WT* and *dfm1Δ* cells treated with 0.4mM of auxin were subjected to immunoprecipitation with HA antibodies. Eluted proteins were analyzed by SDS-PAGE followed by Mass Spectrometry. (D) Degradation of iRC and its variant iRC(Pro) with three point mutations that abolish the tight folding of the DHFR domain in *WT* and *dfm1Δ* cells. Degradation of iRC and iRC(Pro) was induced at time 0 with 0.4mM auxin. Samples were prepared and analyzed as in (A).

### iRC DHFR domain accumulates in Dfm1 mutant cells

Derlins are small multi-pass ER membrane proteins implicated in the degradation of luminal (Knop et al., 1996; Lilley and Ploegh, 2004; Ye et al., 2004; Oda et al., 2006) and membrane (Lilley and Ploegh, 2004; Ye et al., 2004; Stolz et al., 2010; Neal et al., 2018) ERAD substrates. They share similarity with rhomboid intramembrane proteases but lack the catalytic activity (Greenblatt et al., 2011) and there are suggestions that they may be involved in retrotranslocation (Lilley and Ploegh, 2004; Ye et al., 2004; Mehnert et al., 2014; Neal et al., 2018; Wu et al., 2020). We tested the role of the yeast derlins Der1 and Dfm1 in iRC retrotranslocation and degradation. In the absence of Der1, Dfm1 or both, the degradation kinetics of full-length iRC appeared unaffected (Figure 2B). Simultaneous deletion of Der1 or Dfm1 with all ERAD ubiquitin ligases also did not affect turnover of full-length iRC. Strikingly, in all cells lacking Dfm1, auxin-dependent degradation of iRC resulted in the prominent accumulation of a 37kDa fragment. This fragment was smaller than the one described earlier and appeared in much higher amounts (Figure 2B, compare # and *). In contrast with other reports (Neal et al., 2018), this phenotype of *dfm1Δ* was very stable and was not suppressed over time, even after high number of cells divisions.

To determine the identity of the fragments generated in *WT* and *dfm1Δ* cells, they were immunoprecipitated and analyzed by mass spectrometry. Both the ∼40 and 37kDa fragments, hereafter called L (long) and S (short) fragments respectively, contain the luminal domains and transmembrane segment of the iRC, varying solely in the size of their cytosolic tails (Figure 2C). While the S fragment contained 16 cytosolic residues adjacent to the TMD, the L fragment included an additional portion of the AID* tag (Figure 2C and Figure S1). Together, these data show that Dfm1 plays a unique and essential role in the degradation of iRC luminal domain containing DHFR.

Next, we asked whether the accumulation of the S fragment was a direct consequence of the tight folding of DHFR. To this end, we took advantage of DHFR(Pro), a mutant DHFR (Ala29Pro, Trp30Pro, Phe31Pro) which has impaired folding stability (Bhamidipati et al., 2005). An iRC derivative with mutant DHFR(Pro), iRC(Pro), was still degraded in an auxin-dependent manner and with comparable kinetics to original iRC both in *WT* and *dfm1Δ* cells. However, in *dfm1Δ* cells iRC(Pro) was degraded entirely, without accumulation of the S fragment suggesting that Dfm1 requirement increases for substrates with a tightly folded domain in the ER lumen (Figure 2D).

### Retrotranslocation of a tightly folded domain requires Dfm1

Next, we investigated the nature of the defect in *dfm1Δ* cells leading to accumulation of the S fragment. Fractionation experiments showed that both L and S fragments, generated in *WT* and *dfm1Δ* cells respectively, partition with cellular membranes (Figure 3A). This is consistent with the presence of iRC transmembrane segment in both fragments.

**Figure 3.**
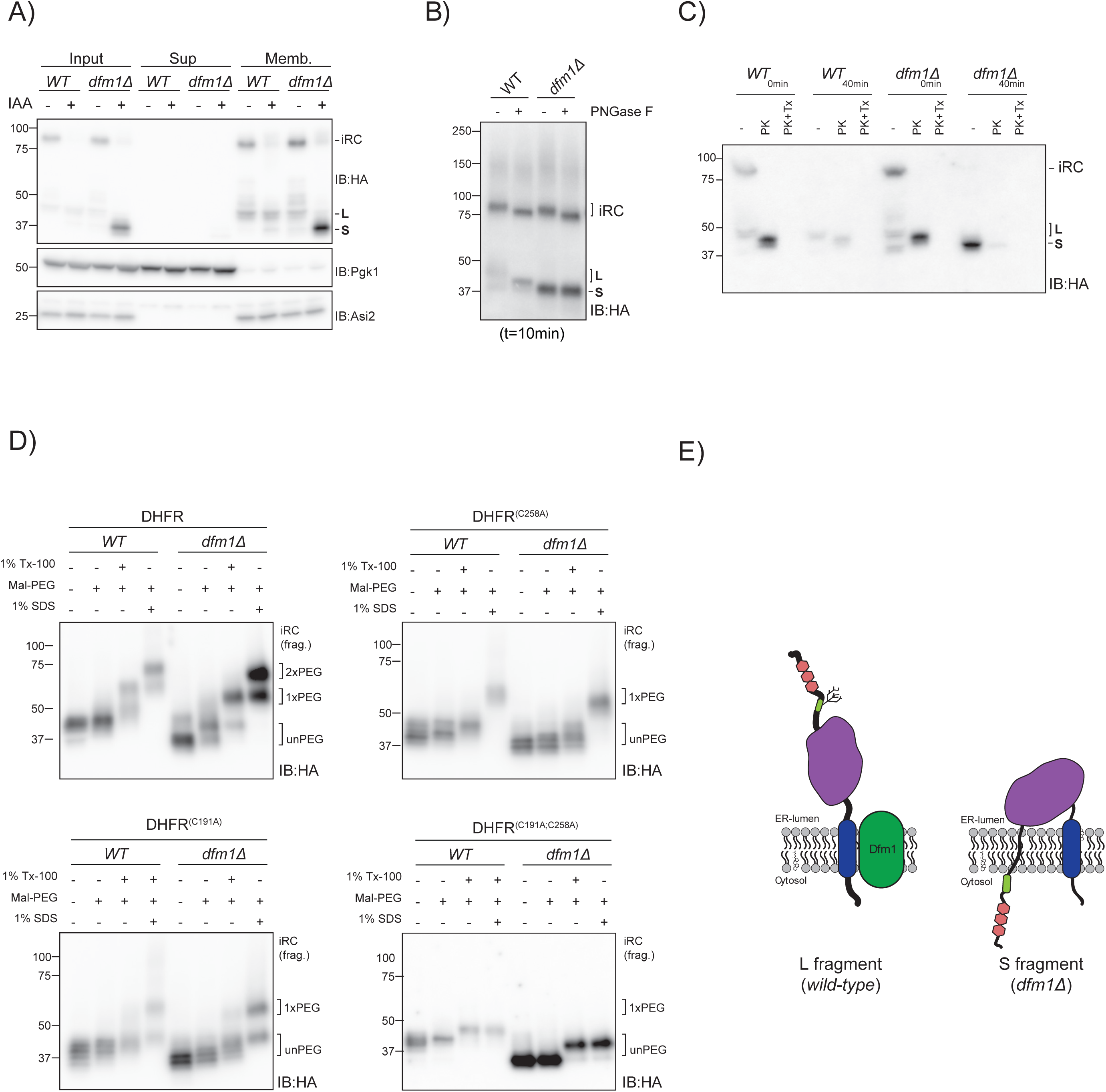
Dfm1 is required for the retrotranslocation of DHFR folded domain from iRC. (A) iRC-derived L and S fragments are membrane-associated. Cell extracts (Input) from *WT* and *dfm1Δ* cells were fractionated in crude membranes (Memb.) and soluble (Sup) material. Where indicated, cells were incubated with IAA (0.4mM for 40 minutes). Fractions were analyzed by SDS-PAGE and western blotting using α-HA antibody. The ER membrane protein Asi2 and the cytosolic soluble Pgk1 were used as controls. (B) Glycosylation status of iRC determined by PNGase F digestion. Denaturing extracts from *WT* and *dfm1Δ* cells incubated with IAA (0.4mM for 10 minutes) were prepared. Upon dilution, extracts were treated with PNGase F and analyzed by SDS–PAGE and western blotting using α-HA antibody. (C) Accessibility of iRC and derived fragments to exogenous proteinase K. ER-derived microsomes from untreated or IAA-treated (0.4mM for 40 minutes) *WT* and *dfm1Δ* cells were incubated with Proteinase-K (PK) (50μg/mL) in the presence or absence of 1% Triton X-100 (Tx). Samples were analyzed by SDS-PAGE followed by western blotting using α-HA antibody. (D) Accessibility of iRC and derived fragments to the cysteine-modifying reagent Maleimide-PEG^5000^ (Mal-PEG). Native extracts from IAA-treated (0.4mM for 40 minutes) *WT* and *dfm1Δ* cells were incubated with 5 mM of Mal-PEG in the absence or presence of the detergents Triton X-100 (Tx-100) or SDS, as indicated. Reactions were subjected to immunoprecipitation with HA antibody, and bound proteins were analyzed by SDS-PAGE and immunoblotting with an α-HA. Note that iRC-derived fragments contain two cysteines, both in the DHFR domain (C191 and C258). The number of cysteine residues modified with Mal-PEG (0, 1 or 2) are indicated by unpeg, 1xPEG and 2xPEG. (E) Diagram representation of the topology adopted by the iRC-derived L and S fragments in *WT* and *dfm1Δ* cells, respectively.

To examine the topology of iRC fragments in relation to the ER membrane we first analyzed their glycosylation status by treatment with the Peptide:N-glycosidase F (PNGase F) enzyme. To generate iRC fragments, *WT* and *dfm1Δ* cells were incubated with auxin for 10 minutes. Subsequently, whole cell lysates from these cells were treated with PNGase F and analyzed by SDS-PAGE and western blotting. In *WT* and *dfm1Δ* cells, PNGase F treatment resulted in faster migration of full-length iRC, as expected (Figure 3B). Similarly, PNGase F digestion of *WT* extracts resulted in faster migration of the L fragment indicating that the N-terminal glycan was in the lumen of the ER, protected from cytosolic peptide N-glycanase. In contrast, PNGase F treatment of extracts from *dfm1Δ* cells did not affect the mobility of the S fragment suggesting that the N-glycan proximal to the N-terminus had been exposed to and processed by endogenous peptide N-glycanase in the cytoplasm (Figure 3B).

To independently validate the topology of auxin-induced L and S fragments, we performed protease protection assays. In the absence of auxin, proteinase-K treatment of microsomes prepared from *WT* and *dfm1Δ* cells showed a similar profile, resulting in a protected fragment of almost 40kDa corresponding to iRC luminal domain (Figure 3C). This data suggest that biogenesis iRC is unaffected in *dfm1Δ* mutant and like in *WT* cells, it has the N-terminus in the ER lumen. A protected fragment of the same size was obtained upon proteinase-K treatment of microsomes prepared from *WT* cells incubated with auxin for 40 minutes. In contrast, proteinase-K treatment of microsomes prepared from *dfm1Δ* cells upon 40-minute auxin incubation did not immunoreact with anti-HA antibodies indicating proteinase-K digestion of the HA epitope at the iRC N-terminus (Figure 3C). Altogether, these data show that in *dfm1Δ* mutants the N-terminus of the S fragment encompassing the HA tag and the N-linked glycan is exposed to the cytosolic leaflet of the ER membrane.

Protease protection and PNGase F assays in WT cells and the experiments with iRC(Pro) suggest that retrotranslocation of tightly folded DHFR challenges efficient iRC degradation. On the other hand, the protease protection and PNGase F results indicate that the N-terminus of iRC adjacent of DHFR accesses the cytosol in *dfm1Δ* cells.

To reconcile these observations and more directly ascertain if the DHFR domain of iRC is effectively retrotranslocated we employed a PEGylation assay with a membrane-impermeable cysteine-reactive reagent, maleimide polyethylene-glycol (mPEG). The iRC-derived fragments generated upon auxin-triggered degradation contain two cysteine residues, both within the DHFR domain. Yeast microsomes prepared from auxin-treated WT and *dfm1Δ* cells were incubated with mPEG in the presence or absence of detergent. After quenching, the samples were immunoprecipitated with an anti-HA antibody and the eluates analyzed by SDS-PAGE and western blotting. In absence of detergent, the fragments remained largely unmodified by mPEG, both in WT and *dfm1Δ* microsomes (Figure 3D). Dissolution of the microsomal membrane with the detergent Triton X-100 resulted in slower migration of the fragments consistent with PEGylation of one of the cysteines (Figure 3D, 1xPEG). This result suggests that the second cysteine is inaccessible to mPEG likely due to sterical constrains in folded DHFR. Indeed, prominent modification of the second cysteine residue was observed in buffer containing 1% SDS, a condition inducing DHFR unfolding (Figure 3D, 2xPEG). These results were confirmed by PEGylation experiments in cells expressing iRC-derivatives with mutations in the two cysteine residues within DHFR (C191A and C258A) (Figure 3D). Together, these experiments show that iRC fragments are in the lumen of the ER. Moreover, the strong accumulation of DHFR in the ER lumen (in the form of S fragment) in the *dfm1Δ* mutant reflects a defect of these cells in the retrotranslocation of tightly folded domains. Of note, the S fragment accumulated in *dfm1Δ* cells displays a unique topological orientation, with its N-terminus facing the cytosol (Figure 3E).

### Distinct Dfm1 domains involved in the retrotranslocation of a tightly folded domain

Dfm1 consists of a rhomboid-like membrane domain followed by an extended cytosolic C-terminus encompassing two SHP boxes shown to bind to Cdc48 (Sato and Hampton, 2006; Greenblatt et al., 2011). Rhomboid-like signature motifs include the WR residues in a luminal loop and the GxxxG sequence in the sixth transmembrane segment (Lemberg and Freeman, 2007). Both WR and GxxxG motifs are required for the function of rhomboid proteases as well as catalytically inactive derlins, such as Dfm1 (Lemberg et al., 2005; Greenblatt et al., 2011; Neal et al., 2018). To test the contribution of Dfm1 rhomboid-like domain in DHFR retrotranslocation we overexpressed the mutants Dfm1^(WR-AA)^ or Dfm1^(GxxxG-AA)^ in *dfm1Δ* cells. While overexpression of Dfm1 prevented accumulation of iRC S fragment, this fragment was still present in cells expressing comparable levels of Dfm1^(WR-AA)^ or Dfm1^(GxxxG-AA)^ (Figure 4A). Curiously, overexpression of Dfm1^(WR-AA)^ in *WT* cells also led to S fragment accumulation indicating a dominant-negative function of this mutation (Figure 4B). Thus, Dfm1 rhomboid-like activity is required for the retrotranslocation and degradation of iRC folded luminal domain.

**Figure 4.**
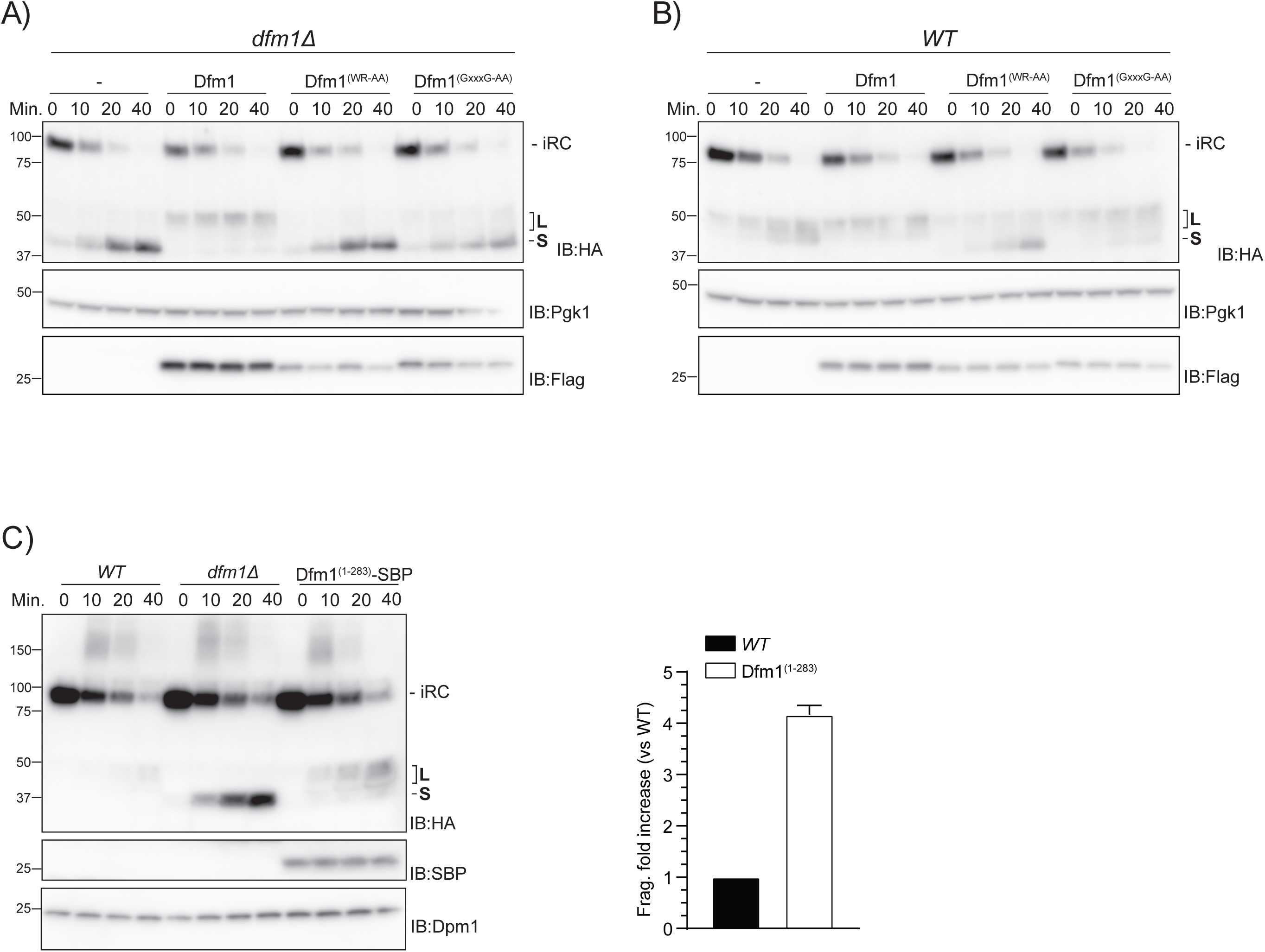
Distinct roles of Dfm1 rhomboid-like motif and Cdc48-interacting domain in iRC retrotranslocation. (A) Time-course analysis of iRC degradation in *dfm1Δ* cells with an empty vector (-) or the indicated *DFM1* variants. Expression of Dfm1 variants was driven from the inducible GAL promoter upon growth in galactose containing media. Cells were treated with 0.4 mM Auxin and cycloheximide at time 0. Whole-cell extracts were analyzed by SDS– PAGE and western blotting. iRC and Flag-tagged Dfm1 derivatives were detected with α-HA and α-Flag antibodies respectively. Pgk1 was used as loading control and detected with α-Pgk1 antibody (B) Time-course analysis of iRC degradation in *WT* cells with an empty vector (-) or overexpressing the indicated *DFM1* variants. Samples were prepared and analyzed as in (A). (C) Time-course analysis of iRC degradation in cells with the indicated genotype. The Dfm1(1-283)-SBP lacks the SHP-motifs shown to interact with the Cdc48 ATPase. Samples were prepared and analyzed as in (A). On the right, the graph shows the amount of L fragment in Dfm1(1-283)-SBP cells in relation to the *WT* at the 40 minutes time point. At least three independent experiments were quantified. Error bars indicate the standard error of the mean.

To examine the role of Dfm1 SHP boxes in iRC degradation we generated Dfm1^(1-283)^, a C-terminal truncation that removes both SHP motifs. Dfm1^(1-283)^ did not affect the degradation rate of the full-length iRC however it led to a four-fold increase in the accumulation of L fragment (Figure 4C). These data suggest that Dfm1 rhomboid-like signature motifs and Cdc48-interacting SHP boxes have distinct roles in retrotranslocation of iRC folded domain.

### Dfm1 and the Cdc48 complex cooperate in the retrotranslocation of a luminal folded domain

The cytosolic Cdc48 ATPase, with the co-factors Npl4 and Ufd1, plays a central role in protein retrotranslocation from the ER (Wu and Rapoport, 2018). To test the role of Cdc48 complex in iRC retrotranslocation and degradation, we used a well-characterized temperature-sensitive (ts) allele, *NPL4-1*, which can be inactivated at 37°C (Ye et al., 2001; Hitchcock et al., 2003). At this temperature atTir1 is non-functional (Nishimura et al., 2009) and the Tir1 from *Oryza sativa* (OsTir1) was used instead. While OsTir1 is active at a wider temperature range, its auxin-dependent recognition of iRC is less stringent and basal degradation of iRC is observed even in the absence of auxin (Figure S2). At 25°C, the permissive temperature at which *NPL4-1* retains some residual activity, auxin treatment resulted in iRC degradation with the generation of the expected L and S fragments in cells with and without *DFM1*, respectively (Fig5A, left panel). Inactivation of Npl4 by a temperature shift to 37°C led to an increase in the amount of L fragment in comparison to *WT* cells (Fig5A, right panel). Importantly, Npl4 inactivation in *dfm1Δ* cells also resulted in a strong accumulation of L fragment with a concomitant decrease in S fragment. Similar results were obtained upon inactivation of Cdc48 complex using a different allele, *cdc48-6* (Schuberth and Buchberger, 2005; Ruggiano et al., 2016) (Figure 5B). These experiments indicate that Cdc48 complex plays an important role in iRC retrotranslocation and that its function precedes the one of Dfm1.

**Figure 5.**
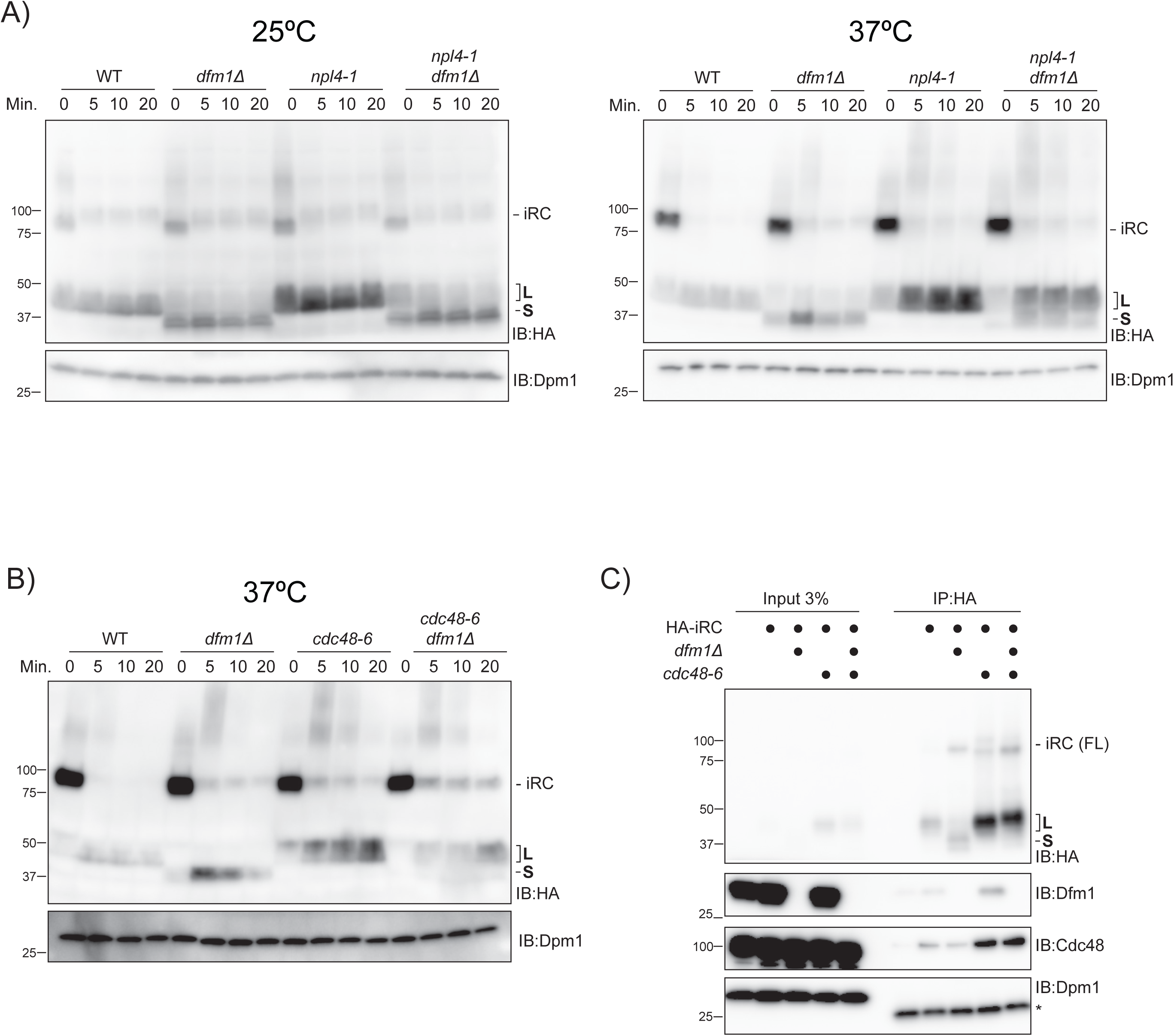
Dfm1 and the Cdc48 complex partner in the retrotranslocation of a luminal folded domain. (A) Time-course analysis of iRC degradation in cells with the indicated genotype at 25°C (left panel) or upon a 60 min shift to 37°C (right panel). At time 0, cells were treated with 50μM Auxin and cycloheximide. In these cells, iRC ubiquitination is mediated by OsTir1, which is functional at 37°C (Nishimura et al., 2009). Whole-cell extracts were analyzed by SDS–PAGE followed by western blotting with α-HA antibody. Dpm1 was used as loading control and detected with α-Dpm1 antibody. (B) Time course analysis of iRC degradation in cells with the indicated genotype, upon a 2h30 shift to 37°C. Samples were prepared and analyzed as in (A). (C) iRC interacts with Dfm1 and Cdc48 ATPase. Cells with the indicated genotype were shifted to 37°C for 2h30 followed by a 20min incubation with auxin (50 μM). Detergent solubilized membranes were subjected to immunoprecipitation with HA antibody, and associated proteins were analyzed by SDS-PAGE and western blotting. HA-tagged iRC was detected with the α-HA antibody. Cdc48, Dfm1 and Dpm1 proteins were detected with indicated antibodies. For the α-Dpm1 blot, the asterisk indicates the light chain of the antibody used for immunoprecipitation.

To investigate the association of iRC with Dfm1 and the Cdc48 complex we used immunoprecipitation. Cells were treated with auxin to trigger iRC degradation and iRC was immunoprecipitated from detergent-solubilized crude membrane fractions. In WT cells, a small fraction of both Dfm1 and Cdc48 associated with iRC. This association increased in cells expressing Cdc48-6, a substrate-trapping allele, suggesting that the interaction of iRC with these factors is transient (Schuberth and Buchberger, 2005; Ruggiano et al., 2016). In *dfm1Δ* cells, iRC interaction with Cdc48-6 was maintained while its association with *WT* Cdc48 appeared slightly reduced. In all cases, the abundant ER membrane protein Dpm1 did not associate with iRC suggesting that its interactions with Dfm1 and Cdc48 were specific.

### Cdc48 recruitment to retrotranslocation intermediates requires Ufd2 and Rad23/Dsk2

ERAD involves various adaptor proteins which facilitate the interaction of substrates with key effectors such as the Cdc48 ATPase or the proteasome (Jentsch and Rumpf, 2007; Wolf and Stolz, 2012). We wondered whether these adaptors were required for iRC degradation. Among ERAD adaptors are Ubx2, involved in recruiting Cdc48 to the ER membrane (Neuber et al., 2005; Schuberth and Buchberger, 2005), and the redundant substrate shuttling factors Rad23 and Dsk2 (Medicherla et al., 2004; Tsuchiya et al., 2017). The degradation of iRC in *ubx2Δ* cells was indistinguishable from WT cells (Figure S3). In contrast, *rad23Δ dsk2Δ* mutant cells showed strongly impaired iRC degradation and even the generation of L fragment was defective (Figure 6A). Strikingly, a similar phenotype was observed in cells lacking Ufd2, a polyubiquitination enzyme shown to bridge Cdc48 to Rad23 and Dsk2 (Richly et al., 2005; Baek et al., 2011) (Figure 6A). These results suggested that Rad23/Dsk2 and Ufd2 were required upstream of Cdc48, perhaps from a defect in Cdc48 recruitment to iRC. To directly test this possibility, we analyzed the interaction of Cdc48 with iRC before and after inducing its auxin-dependent ubiquitination. In the absence of auxin no interaction between iRC and Cdc48 was observed, as expected (Figure 6B). However, upon iRC ubiquitination by addition of auxin resulted in its interaction with Cdc48. The interaction was specific as the other abundant complexes with ubiquitin-binding activity, such as the proteasome did not co-precipitate with iRC. Dfm1 stabilizes Cdc48 interaction with iRC, as less Cdc48 co-precipitated with iRC in *dfm1Δ* cells. Remarkably, the interaction between iRC and Cdc48 was lost both in *rad23Δdsk2Δ* and in *ufd2Δ* cells (Figure 6B and Figure S4). Together these experiments indicate that Rad23/Dsk2 shuttle factors, together with Ufd2, facilitate the recruitment of Cdc48 to iRC.

**Figure 6.**
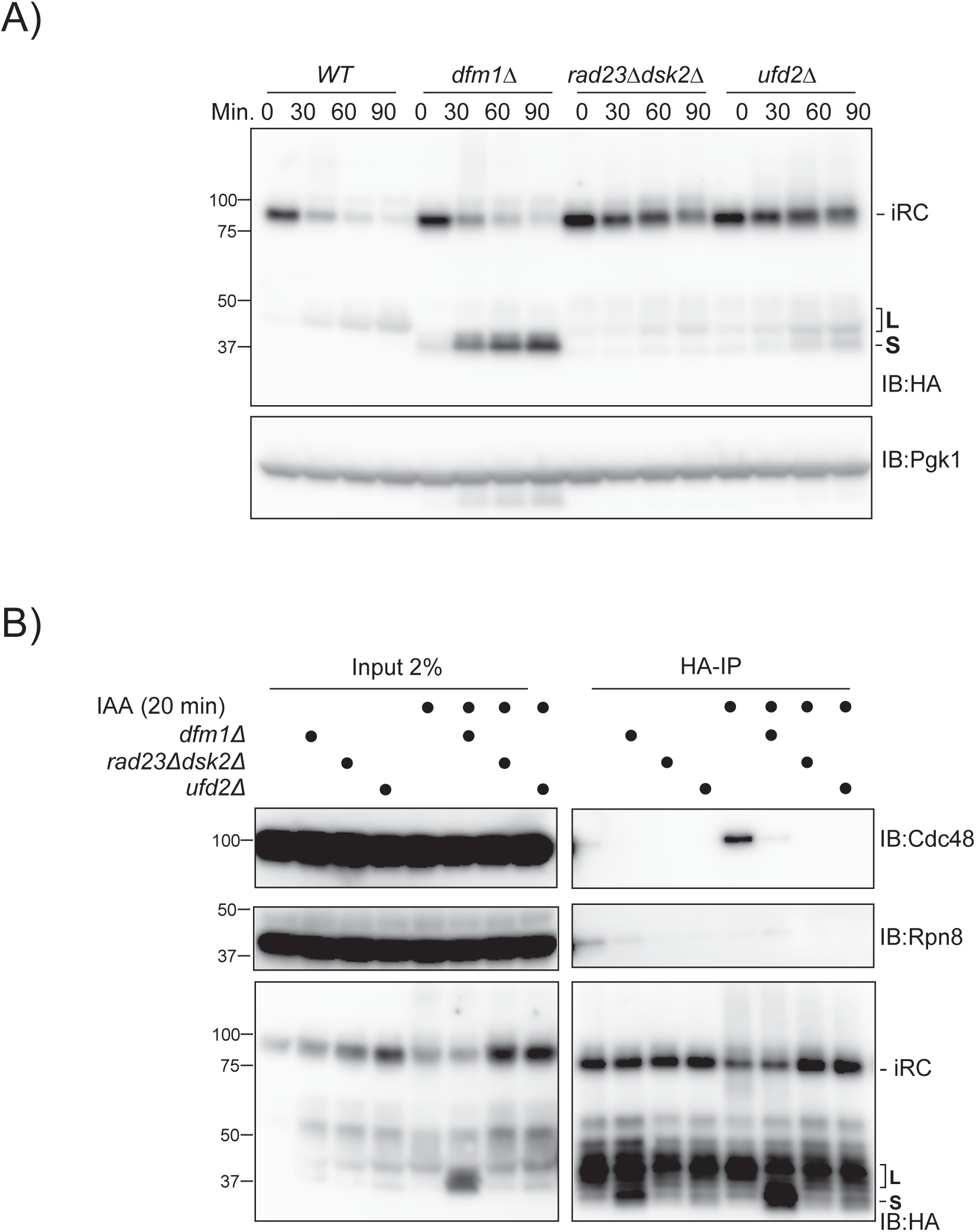
iRC processing by Cdc48 and degradation require Ufd2 and the shuttle factors Rad23/Dsk2. (A) Time-course analysis of iRC degradation in cells with the indicated genotype. Samples were prepared and analyzed as in Figure 1(E). (B) iRC interaction with Cdc48 ATPase depends on its auxin-dependent degradation and is impaired in the indicated mutants. Cells with the indicated genotype were incubated in absence or presence of IAA (20 minutes). Cell extracts were subjected to immunoprecipitation with HA antibody, and associated proteins were analyzed by SDS-PAGE and western blotting. HA-tagged iRC was detected with α-HA antibody. Cdc48 and Rpn8 were detected with anti-Cdc48 and anti-Rpn8 antibodies, respectively. Note that in native extracts the iRC substrate is labile and L fragment is generated even in absence of Auxin, as confirmed by analysis of the same samples under denaturing conditions (see Figure S4).

## DISCUSSION

Retrotranslocation of folded protein domains is challenging and their presence slows down the degradation of ERAD substrates (Tirosh et al., 2003; Bhamidipati et al., 2005; Shi et al., 2019). However, how proteins with folded luminal domains are retrotranslocated across the ER membrane is unknown. To investigate this issue, we developed the iRC, a conditional substrate whose biogenesis and folding can be uncoupled from its degradation, triggered by auxin.

Based on our data, we propose the following model for iRC retrotranslocation and degradation (Figure 7). In the absence of auxin, the iRC is a long-lived ER membrane protein consistent with it being properly folded and not recognized by quality control processes such as ERAD. Auxin-induced iRC ubiquitination results in its conversion into the L fragment as well as in its association with the Cdc48/Npl4/Ufd1 ATPase complex, both requiring the ubiquitin shuttle factors Rad23/Dsk2 and the multiubiquitin chain extension enzyme Ufd2. The proteasome has been implicated in proteolytic clipping of several ubiquitinated ER proteins (Hoppe et al., 2000; Kohlmann et al., 2008) and likely also contributes to the L fragment generation, a possibility should be directly tested in future studies.

**Figure 7.**
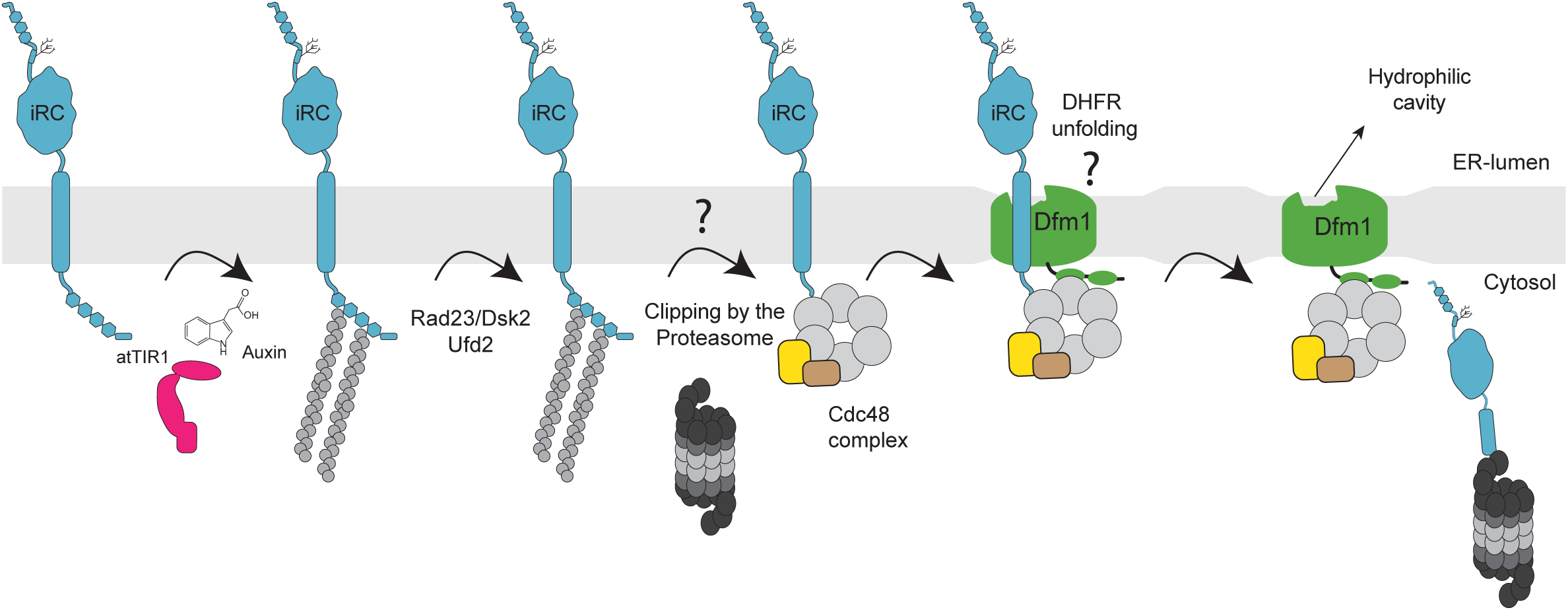
An outline of the pathway for iRC degradation. See discussion for details.

The role of Rad23/Dsk2 and Ufd2 in Cdc48 complex recruitment should be also further characterized. These factors shuttle ubiquitinated substrates from Cdc48 to the proteasome (Medicherla et al., 2004; Richly et al., 2005; Baek et al., 2011). However, recent studies in human cells demonstrated that RAD23B, the human homolog of Rad23, can also function in concentrating ubiquitinated substrates for subsequent processing by p97, the Cdc48 homolog, and the proteasome (Yasuda et al., 2020). Interestingly, this role of RAD23B was observed specifically during hyperosmotic stress, a condition that like auxin-induced ubiquitination of iRC, leads to an acute increase in ubiquitinated substrates. Dfm1 has also been shown to facilitate Cdc48 association with ER membranes. (Neal et al., 2018). Consistently, while Cdc48 can be recruited to iRC in *dfm1Δ* cells, persistent association of Cdc48 with the iRC requires Dfm1, in particular its SHP boxes. Indeed, cells expressing Dfm1(1-283), a truncated Dfm1 lacking the SHP boxes, showed increased accumulation L fragment, just like Cdc48 complex mutants. Thus, Rad23/Dsk2, Ufd2 and Dfm1 all appear to contribute to the stable engagement of Cdc48 complex with iRC. The Cdc48 complex, through rounds of ATP hydrolysis, pulls on iRC domain exposed to the cytoplasm while Dfm1, through its rhomboid-like membrane region, interacts with and facilitates the retrotranslocation of iRC folded luminal domain. The joint activity of Cdc48 complex and Dfm1 results in iRC release from the membrane for subsequent degradation by the proteasome (Figure 7).

Dfm1 was necessary for the degradation of iRC while iRC(Pro), an analogous substrate containing a luminal unfolded DHFR domain, was degraded independently of Dfm1. The specific requirement of Dfm1 for substrates with folded luminal domains suggests that it has a role in the resolution of stalled retrotranslocation intermediates. Whether this is achieved by promoting the retrotranslocation of luminal domains in their folded state or by assisting their unfolding prior retrotranslocation is an important open question.

In the absence of Dfm1, iRC accumulates in the form of S fragment. This fragment, which is generated in a Cdc48-dependent manner, has an interesting topology, with the N-terminus exposed to the cytoplasm while the folded DHFR domain fails to retrotranslocate and remains within the ER lumen (Figure 3E). Whether S fragment is a general intermediate in iRC degradation or an off-pathway product generated only in the absence of Dfm1 is unclear.

Dfm1 rhomboid-like signature motifs are required for the retrotranslocation of folded DHFR domain as observed by the accumulation of S fragment in cells expressing Dfm1 rhomboid-like signature mutants. In a recent cryo-EM structure of the Dfm1 homolog Der1, these rhomboid-like signature motif form a hydrophilic cavity on the luminal side of the ER membrane (Wu et al., 2020). As observed in previous structures of active rhomboid proteases (Wang et al., 2006; Wu et al., 2006; Ben-Shem et al., 2007), the signature motif also leads to thinning of the surrounding membrane. In principle, both the hydrophilic cavity and the ability to thin the membrane may facilitate Dfm1-mediated retrotranslocation.

Importantly, expression of Dfm1 rhomboid-like signature motif mutants leads to S fragment accumulation even in cells with *WT* Dfm1. The dominant-negative effect of these mutants suggests that Dfm1 functions as an oligomer, as previously shown by immunoprecipitation experiments (Goder et al., 2008; Stolz et al., 2010). Alternatively, mutant Dfm1 may titrate away an essential component for iRC degradation, such as Cdc48 or other currently unknown factors. While this can only be clarified experimentally, it should be mentioned that rhomboid proteases notably function as monomers (Kreutzberger and Urban, 2018). In addition, a cryo-EM structure of the Hrd1 complex required for the retrotranslocation of luminal misfolded proteins contains a single Der1 molecule (Wu et al., 2020).

Dfm1 was recently implicated in the retrotranslocation of misfolded membrane proteins (Stolz et al., 2010; Neal et al., 2018). These include substrates of both Doa10- and Hrd1-dependent ERAD, consistent with previous immunoprecipitation studies which showed interactions between Dfm1 and the two ubiquitin ligases (Goder et al., 2008; Stolz et al., 2010). Curiously, the requirement for Dfm1 in retrotranslocation of misfolded membrane proteins is transient. While these substrates are stabilized in freshly generated *dfm1Δ* cells, after a few generations these mutants are capable of degrading misfolded membrane proteins with a kinetics similar to *WT* cells (Neal et al., 2018). The spontaneous and rapid phenotypic suppression in *dfm1Δ* involved the duplication of the *HRD1* locus pointing to redundant roles of Hrd1 and Dfm1 in the degradation of misfolded membrane proteins. In contrast, we observe that the requirement of Dfm1 in the degradation of iRC degradation is stable over time and was independent of *HRD1* as well as other ERAD components. Despite the differences, the retrotranslocation of both misfolded and folded membrane proteins involves rhomboid-like signature motifs of Dfm1 as well as its SHP boxes for Cdc48 interaction. Thus, the degradation of misfolded and folded membrane proteins appear to have common as well as distinct steps, both involving Dfm1. It will be interesting to directly compare the roles of Dfm1 in the retrotranslocation of the two types of substrates.

Here, we used a single substrate, the iRC, to delineate a simple pathway for the retrotranslocation of membrane proteins with folded luminal domains. While the iRC will be useful for mechanistic studies, it will be of great interest to identify additional substrates following the same degradation route. Several membrane proteins are targeted for degradation upon ubiquitination by soluble ubiquitin ligases. This group include for example SUN2 (Coyaud et al., 2015) or its yeast homologue Mps3 (Koch et al., 2019), both with large folded domains in the ER lumen. Importantly, in neither case, the membrane components required for the degradation have been identified. Our work indicates that Dfm1 or mammalian Derlins are good candidates to test.

## METHODS

### Yeast strains and Plasmids

The strains used are isogenic either to BY4741 (MATa *ura3Δ0 his3Δ1 leu2Δ0 met15Δ0*) or FY251 (*MATa ura3-52 his3Δ200 leu2Δ1 trp1Δ63*) and are listed in the Supplementary Table S1. Tagging of proteins and individual gene deletions were performed by standard PCR-based homologous recombination (Longtine et al., 1998) or CRISPR (Laughery et al., 2015). The Dfm1(1-283-SBP) allele lacking residues (284-341) with SBP-tag was generated by CRISPR-based gene editing as described (Laughery et al., 2015). In brief, a single guide RNA sequence targeting the desired region of Dfm1 was designed using online software (http://wyrickbioinfo2.smb.wsu.edu) and cloned into a pML vector (Addgene) containing the Cas9 endonuclease from Streptococcus pyogenes using restriction ligation. Cas9 plasmid along with a PCR amplified template containing the deletion of AA (284-341) were transformed using standard transformation protocol. Colonies were screened by PCR and sequencing of genomic DNA. Positive clones were grown in rich media (YPD) for 2-3 days to allow the loss of Cas9 plasmid.

Strains with multiple gene deletions and temperature-sensitive alleles were made by crossing haploid cells of opposite mating types, followed by sporulation and tetrad dissection using standard protocols (Fink and Guthrie, 1991). All cloning procedures were carried out with PCR and Gibson-based assembly or restriction ligation. Dfm1-FLAG construct was generated by genomic amplification of *DFM1* gene and cloned into a pRS425-Gal with a single FLAG tag at the C-terminus. Dfm1 signature motif mutants were generated by sit-directed mutagenesis. Coding sequences from iRC construct and O. sativa *TIR1* were ordered as gene blocks. For all strains, a minimal domain of IAA17 consisting of amino acids 71-114 as described in (Morawska and Ulrich, 2013) was used. Following its verification by sequencing, the iRC and variant constructs were amplified inserted either with a CPY promoter or an ADH promoter into the *HO* locus of all strains of interest. Plasmids used in this study are listed in Supplementary Tables S2.

### Substrate degradation experiments

Cycloheximide shutoff chases were performed as described previously (Foresti et al., 2014.) Briefly, yeast cells were grown either in rich media (YPD) or synthetic media with 2% glucose in exponential phase (OD_600_ 0.6 - 0.9) at 25°C. Over-expression of Dfm1-FLAG variants was driven from the inducible GAL promoter upon constant growth synthetic media (SC-Leucine) with 4% D-Galatactose to exponential phase at 25°C. 6 ODs of cells were harvested, centrifuged for 2 min at 3500rpm and resuspended in the appropriate media at 1 OD/mL. For auxin-dependent degradation experiments, auxin (Indole-3-acetic acid, IAA) was added to the cultures from a stock prepared in DMSO to a final concentration of 400 μM. For experiments using OsTir1, cells were treated with 50 μM of auxin. In experiments with cycloheximide, the drug was used at 250 μg/mL from a stock of 12.5 mg/mL prepared in H_2_O. At the specified time points 1 OD of cells was collected and whole-cell lysates were prepared using NaOH extraction - cell pellets were treated with 300 μL of 150 mM NaOH for 10 min on ice, briefly centrifuged and resuspended in 50 μL sample buffer (SB; 100 mM Tris/HCl [pH6.8], 3% SDS, 15% glycerol, 75 mM DTT). Samples were then analyzed by SDS-PAGE (0.4 ODs per sample were loaded) followed by immunoblot with the indicated antibodies. Unless indicated, chases were performed at 25°C. Temperature-sensitive mutants were grown at 25°C and shifted to 37°C for 1-2.5 hours prior to the addition of auxin. For the experiments with galactose induction, cells were grown directly at 25°C in SC media supplemented with 4% galactose. Data quantification was performed using Image Studio software (Li-Cor) and graphs were plotted in GraphPad Prism v6. Representative images of three independent experiment are shown.

### Deglycosylation experiments

Cell lysates obtained by NaOH extraction were diluted 1:1 in water to final concentration of SB (50mM Tris/HCl [pH6.8], 1.5% SDS, 7.5% glycerol, 37.5mM DTT). PNGaseF (0.5 μL, 2500U) of was added to 20 μL of lysate and incubated for 15 min at 37°C. Samples were then analyzed by SDS-PAGE followed by immunoblot.

### Membrane fractionation experiments

Cells fractionation experiments 50 ODs of cells were lysed in LB buffer (50mM Tris/HCl [pH7.4], 200 mM NaCl, 1mM EDTA, 2mM phenylmethylsulfonyl fluoride (PMSF) and protease inhibitor cocktail) by 5-6 x 1 min cycles of bead-beating at 4*C. Lysates were cleared by a 10 min centrifugation at 2000 rpm. Crude membrane and soluble fractions were obtained from cleared lysates by centrifugation at 45000 rpm (25 min at 4°C), using a TLA 100.3 rotor. The membrane pellet was resuspended in sample buffer (100 mM Tris/HCl [pH6.8], 3% SDS, 15% glycerol, 75 mM DTT) and solubilized at 65°C for 10 min.

### Co-Immunoprecipitation

100 ODs of cells were harvested and lysed as described for membrane fractionation experiments, except that the membrane fraction was solubilized in LB + 1% GDN (glyco-diosgenin) supplemented with 1mM PMSF and protease inhibitor cocktail. After 2 hours, the solubilized membranes were precleared by centrifugation for 15 min at max speed on a table-top centrifuge and the supernatant was incubated overnight with 20 μL of anti-HA magnetic beads (Pierce TM). Beads were washed 3 times with LB + 0.02% GDN and bound proteins eluted with SB buffer and analyzed by SDS-PAGE and immunoblotting.

### Protease protection assay

50 ODs of cells were harvested and lysed as described for membrane fractionation experiments, except that the membrane fraction was resuspended in buffer A (20 mM HEPES [pH7.4], 2 mM magnesium acetate, 150 mM KCl, 200 mM sucrose) that lacked protease inhibitors. Total lysates were incubated with Proteinase K (50 μg/mL) or Proteinase K with 1% Triton X-100 for 30 min on ice. The reaction was stopped by the addition of PMSF (2.5 mM final concentration) for 10 min on ice, and after the reaction products were diluted 1:1 in SB and analyzed by SDS–PAGE and immunoblotting.

### PEGylation assays

Cells were grown in YPD until OD of 0.75 and added 0.5 mM auxin for 40min. 250 ODs of cells were harvested and treated with 10 mM Tris-HCl [pH10.5], 1 mM DTT for 10min at room temperature. Cells were washed with SP buffer (50 mM Tris/HCl pH 7.5, 100 mM NaCl, 0.3 M Sorbitol, 1 mM EDTA, 1 mM phenylmethylsulphonyl fluoride (PMSF), 2 mM benzamidine and cOmplete protease inhibitor cocktail (Roche Diagnostics, Mannheim, Germany)) and incubated with zymolyase 20T for ∼20min at 37°C. Spheroplasts were washed, resuspended in 5 mL of SP buffer, and were dounced to generate microsomes. Microsome fraction was centrifugated for 5min at 2000rpm to remove unbroken cells. Microsomal lysate (300 μL) were treated with 2.5 mM of methoxy-polyethylene glycol maleimide (Mal-PEG 5K) for 1h on ice and the reaction was quenched with 25 mM DTT for 30 min. In detergent treated reactions, microsomes were mixed with 1% Triton X-100 or 1% SDS prior to the addition of Mal-PEG in a total volume reaction of 400μL. Samples were diluted to 1.2mL in LB supplemented with 1% Triton X-100 and incubated overnight with 20 μL anti-HA beads (Pierce TM). Beads were washed 3 times with LB + 1% Triton X-100 and bound proteins eluted with 20 μL SB buffer and analyzed by SDS-PAGE and immunoblotting.

### Mass spectrometry

Cells were grown in YPD until OD of 0.75 and added 0.5 mM auxin for 40min. 150 ODs of cells were harvested, and membrane fraction was prepared using bead beating as described above. The pelleted membranes were solubilized in 1.2mL in LB supplemented with 1% Triton X-100 and incubated overnight with 75 μL anti-HA beads (Pierce TM). Beads were washed 3 times with LB +1% Triton X-100 and bound proteins eluted with 100 μL SB buffer and 95% was separated by SDS-PAGE and Coomassie-stained. In parallel immunoblotting was performed in 5% of the same sample against HA epitope. The fragments bands were excised from the Coomassie-stained gel and subjected to in-gel trypsin digestion as previously described (Shevchenko et al., 2006). After digestion, peptides were separated by nano-flow reversed-phase liquid chromatography coupled to Q Exactive Hybrid Quadrupole-Orbitrap mass spectrometer (Thermo Fisher Scientific). Peptides were loaded on a C18 PepMap100 pre-column (inner diameter 300μm×5mm, 3 μm C18 beads; Thermo Fisher Scientific) and separated on a 50 cm reversed-phase C18 column (inner diameter 75 μm, 2 μm C18 beads) using a linear gradient from 10 to 35% of B for 30 min at a flow rate of 200 nL/min (A: 0.1% formic acid, B: 0.1% formic acid in acetonitrile). All data were acquired in a data-dependent mode, automatically switching from MS to collision-induced dissociation MS/MS on the top 10 most abundant ions with a precursor scan range of 350–2000m/z. MS spectra were acquired at a resolution of 70 000 and MS/MS scans at 17 500. Dynamic exclusion was enabled with an exclusion duration of 5 s and charge exclusion was applied to unassigned, and mono-charged ions. Raw data files were processed for protein identification using MaxQuant, version 1.6.3.4 (Tyanova et al., 2016). The MS/MS spectra were searched against the relevant Uniprot proteome database; precursor mass tolerance was set to 20 ppm and MS/MS tolerance to 0.05 Da. Enzyme specificity was set to trypsin with a maximum of two missed cleavages. False discovery rate for protein and peptide spectral matches was set at 0.01.

## Acknowledgements

We thank M. Glickman, F. Wilfing and D Wolf for reagents, members of the Carvalho lab for discussions and M Freeman, L. Krshnan, T. Rapoport, J. Robson-Tull and M. Weijer for comments on the manuscript. Proteomics experiments were performed at Advanced proteomics facility, Department of Biochemistry, University of Oxford. P. Carvalho was supported by an investigator award from The Wellcome Trust (202642/Z/16/Z).

## Legends to the Supplemental Figures

**Figure S1. Mass spectrometry analysis of iRC-derived L and S fragments.**

(A) Amino acid sequence of iRC. Underlined in blue and red are the peptides identified by mass spectrometry for the L and S fragments, respectively.

(B) The intensity of the peptides from the of L and S fragments in *WT* and *dfm1Δ* cells, respectively, as determined by mass spectrometry.

**Figure S2. Auxin-independent degradation iRC in cells expressing Oryza sativa Tir1 (OsTir1).**

*WT* cells expressing the iRC with and without V5-tagged OsTir1 (V5-OsTir1) were grown in YPD media in the absence of auxin at the indicated temperature. Whole cell lysates were analyzed by SDS-PAGE and immunoblotting with the indicated antibodies. Dpm1 was used as loading control and detected with α-Dpm1 antibody.

**Figure S3. Mutation of the Cdc48 adaptor Ubx2 does not affect iRC retrotranslocation and degradation.**

The degradation of iRC in cells with the indicated genotype was analyzed as in Figure 1. In this experiment, iRC was expressed from the strong constitutive *ADH* promoter resulting in > 10 fold higher steady state levels of iRC, in comparison to a iRC construct expressed from the weaker *CPY* promoter, used in the majority of the experiments. The genetic requirements for iRC turnover of were the same irrespective of the promoter used. However, when expressed from the strong *ADH* promoter, the turnover of iRC was slower and resulted in higher amounts of L and S fragments in WT and *dfm1Δ* cells, respectively. This is consistent with the retrotranslocation of DHFR being rate-limiting.

**Figure S4. Samples used in Figure 6B analyzed under denaturing conditions.** Whole cells extracts prepared under denaturing conditions were analyzed by SDS-PAGE and immunoblotting.

## References

Arkowitz, R.A., Joly, J.C., and Wickner, W. (1993). Translocation can drive the unfolding of a preprotein domain. EMBO J. 12, 243–253.

Baek, G.H., Kim, I., and Rao, H. (2011). The Cdc48 ATPase modulates the interaction between two proteolytic factors Ufd2 and Rad23. Proc. Natl. Acad. Sci. USA 108, 13558–13563.

Baldridge, R.D., and Rapoport, T.A. (2016). Autoubiquitination of the hrd1 ligase triggers protein retrotranslocation in ERAD. Cell 166, 394–407.

Bays, N.W., Wilhovsky, S.K., Goradia, A., Hodgkiss-Harlow, K., and Hampton, R.Y. (2001). HRD4/NPL4 is required for the proteasomal processing of ubiquitinated ER proteins. Mol. Biol. Cell 12, 4114–4128.

Ben-Shem, A., Fass, D., and Bibi, E. (2007). Structural basis for intramembrane proteolysis by rhomboid serine proteases. Proc. Natl. Acad. Sci. USA 104, 462–466.

Bhamidipati, A., Denic, V., Quan, E.M., and Weissman, J.S. (2005). Exploration of the topological requirements of ERAD identifies Yos9p as a lectin sensor of misfolded glycoproteins in the ER lumen. Mol. Cell 19, 741–751.

Bhattacharya, A., and Qi, L. (2019). ER-associated degradation in health and disease - from substrate to organism. J. Cell Sci. 132.

Bodnar, N.O., and Rapoport, T.A. (2017). Molecular mechanism of substrate processing by the cdc48 atpase complex. Cell 169, 722–735.e9.

Carvalho, P., Goder, V., and Rapoport, T.A. (2006). Distinct ubiquitin-ligase complexes define convergent pathways for the degradation of ER proteins. Cell 126, 361–373.

Carvalho, P., Stanley, A.M., and Rapoport, T.A. (2010). Retrotranslocation of a misfolded luminal ER protein by the ubiquitin-ligase Hrd1p. Cell 143, 579–591.

Christianson, J.C., and Ye, Y. (2014). Cleaning up in the endoplasmic reticulum: ubiquitin in charge. Nat. Struct. Mol. Biol. 21, 325–335.

Coyaud, E., Mis, M., Laurent, E.M.N., Dunham, W.H., Couzens, A.L., Robitaille, M., Gingras, A.-C., Angers, S., and Raught, B. (2015). BioID-based Identification of Skp Cullin F-box (SCF)β-TrCP1/2 E3 Ligase Substrates. Mol. Cell Proteomics 14, 1781–1795.

Denic, V., Quan, E.M., and Weissman, J.S. (2006). A luminal surveillance complex that selects misfolded glycoproteins for ER-associated degradation. Cell 126, 349–359.

Eilers, M., and Schatz, G. (1986). Binding of a specific ligand inhibits import of a purified precursor protein into mitochondria. Nature 322, 228–232.

Fink, G.R., and Guthrie, C. (1991). Guide to yeast genetics and molecular biology.

Freeman, M. (2014). The rhomboid-like superfamily: molecular mechanisms and biological roles. Annu. Rev. Cell Dev. Biol. 30, 235–254.

Gambill, B.D., Voos, W., Kang, P.J., Miao, B., Langer, T., Craig, E.A., and Pfanner, N. (1993). A dual role for mitochondrial heat shock protein 70 in membrane translocation of preproteins. J. Cell Biol. 123, 109–117.

Goder, V., Carvalho, P., and Rapoport, T.A. (2008). The ER-associated degradation component Der1p and its homolog Dfm1p are contained in complexes with distinct cofactors of the ATPase Cdc48p. FEBS Lett. 582, 1575–1580.

Greenblatt, E.J., Olzmann, J.A., and Kopito, R.R. (2011). Derlin-1 is a rhomboid pseudoprotease required for the dislocation of mutant α-1 antitrypsin from the endoplasmic reticulum. Nat. Struct. Mol. Biol. 18, 1147–1152.

Hitchcock, A.L., Auld, K., Gygi, S.P., and Silver, P.A. (2003). A subset of membrane-associated proteins is ubiquitinated in response to mutations in the endoplasmic reticulum degradation machinery. Proc. Natl. Acad. Sci. USA 100, 12735–12740.

Hoppe, T., Matuschewski, K., Rape, M., Schlenker, S., Ulrich, H.D., and Jentsch, S. (2000). Activation of a membrane-bound transcription factor by regulated ubiquitin/proteasome-dependent processing. Cell 102, 577–586.

Jarosch, E., Taxis, C., Volkwein, C., Bordallo, J., Finley, D., Wolf, D.H., and Sommer, T. (2002). Protein dislocation from the ER requires polyubiquitination and the AAA-ATPase Cdc48. Nat. Cell Biol. 4, 134–139.

Jentsch, S., and Rumpf, S. (2007). Cdc48 (p97): a “molecular gearbox” in the ubiquitin pathway? Trends Biochem. Sci. 32, 6–11.

Knop, M., Finger, A., Braun, T., Hellmuth, K., and Wolf, D.H. (1996). Der1, a novel protein specifically required for endoplasmic reticulum degradation in yeast. EMBO J. 15, 753–763.

Koch, B.A., Jin, H., Tomko, R.J., and Yu, H.-G. (2019). The anaphase-promoting complex regulates the degradation of the inner nuclear membrane protein Mps3. J. Cell Biol. 218, 839–854.

Kohlmann, S., Schäfer, A., and Wolf, D.H. (2008). Ubiquitin ligase Hul5 is required for fragment-specific substrate degradation in endoplasmic reticulum-associated degradation. J. Biol. Chem. 283, 16374–16383.

Kreutzberger, A.J.B., and Urban, S. (2018). Single-Molecule Analyses Reveal Rhomboid Proteins Are Strict and Functional Monomers in the Membrane. Biophys. J. 115, 1755–1761.

Laughery, M.F., Hunter, T., Brown, A., Hoopes, J., Ostbye, T., Shumaker, T., and Wyrick, J.J. (2015). New vectors for simple and streamlined CRISPR-Cas9 genome editing in Saccharomyces cerevisiae. Yeast 32, 711–720.

Lemberg, M.K., and Freeman, M. (2007). Cutting proteins within lipid bilayers: rhomboid structure and mechanism. Mol. Cell 28, 930–940.

Lemberg, M.K., Menendez, J., Misik, A., Garcia, M., Koth, C.M., and Freeman, M. (2005). Mechanism of intramembrane proteolysis investigated with purified rhomboid proteases. EMBO J. 24, 464–472.

Lilley, B.N., and Ploegh, H.L. (2004). A membrane protein required for dislocation of misfolded proteins from the ER. Nature 429, 834–840.

Longtine, M.S., McKenzie, A., Demarini, D.J., Shah, N.G., Wach, A., Brachat, A., Philippsen, P., and Pringle, J.R. (1998). Additional modules for versatile and economical PCR-based gene deletion and modification in Saccharomyces cerevisiae. Yeast 14, 953–961.

Medicherla, B., Kostova, Z., Schaefer, A., and Wolf, D.H. (2004). A genomic screen identifies Dsk2p and Rad23p as essential components of ER-associated degradation. EMBO Rep. 5, 692–697.

Mehnert, M., Sommer, T., and Jarosch, E. (2014). Der1 promotes movement of misfolded proteins through the endoplasmic reticulum membrane. Nat. Cell Biol. 16, 77–86.

Morawska, M., and Ulrich, H.D. (2013). An expanded tool kit for the auxin-inducible degron system in budding yeast. Yeast 30, 341–351.

Natarajan, N., Foresti, O., Wendrich, K., Stein, A., and Carvalho, P. (2020). Quality control of protein complex assembly by a transmembrane recognition factor. Mol. Cell 77, 108–119.e9.

Neal, S., Jaeger, P.A., Duttke, S.H., Benner, C., K Glass, C., Ideker, T., and Hampton, R.Y. (2018). The dfm1 derlin is required for ERAD retrotranslocation of integral membrane proteins. Mol. Cell 69, 306–320.e4.

Needham, P.G., Guerriero, C.J., and Brodsky, J.L. (2019). Chaperoning Endoplasmic Reticulum-Associated Degradation (ERAD) and Protein Conformational Diseases. Cold Spring Harb. Perspect. Biol. 11.

Neuber, O., Jarosch, E., Volkwein, C., Walter, J., and Sommer, T. (2005). Ubx2 links the Cdc48 complex to ER-associated protein degradation. Nat. Cell Biol. 7, 993–998.

Nishimura, K., Fukagawa, T., Takisawa, H., Kakimoto, T., and Kanemaki, M. (2009). An auxin-based degron system for the rapid depletion of proteins in nonplant cells. Nat. Methods 6, 917–922.

Oda, Y., Okada, T., Yoshida, H., Kaufman, R.J., Nagata, K., and Mori, K. (2006). Derlin-2 and Derlin-3 are regulated by the mammalian unfolded protein response and are required for ER-associated degradation. J. Cell Biol. 172, 383–393.

Richly, H., Rape, M., Braun, S., Rumpf, S., Hoege, C., and Jentsch, S. (2005). A series of ubiquitin binding factors connects CDC48/p97 to substrate multiubiquitylation and proteasomal targeting. Cell 120, 73–84.

Ruggiano, A., Foresti, O., and Carvalho, P. (2014). ER-associated degradation: protein quality control and beyond. J. Cell Biol. 204, 869–879.

Ruggiano, A., Mora, G., Buxó, L., and Carvalho, P. (2016). Spatial control of lipid droplet proteins by the ERAD ubiquitin ligase Doa10. EMBO J. 35, 1644–1655.

Sato, B.K., and Hampton, R.Y. (2006). Yeast Derlin Dfm1 interacts with Cdc48 and functions in ER homeostasis. Yeast 23, 1053–1064.

Schoebel, S., Mi, W., Stein, A., Ovchinnikov, S., Pavlovicz, R., DiMaio, F., Baker, D., Chambers, M.G., Su, H., Li, D., et al. (2017). Cryo-EM structure of the protein-conducting ERAD channel Hrd1 in complex with Hrd3. Nature 548, 352–355.

Schuberth, C., and Buchberger, A. (2005). Membrane-bound Ubx2 recruits Cdc48 to ubiquitin ligases and their substrates to ensure efficient ER-associated protein degradation. Nat. Cell Biol. 7, 999–1006.

Shevchenko, A., Tomas, H., Havlis, J., Olsen, J.V., and Mann, M. (2006). In-gel digestion for mass spectrometric characterization of proteins and proteomes. Nat. Protoc. 1, 2856–2860.

Shi, J., Hu, X., Guo, Y., Wang, L., Ji, J., Li, J., and Zhang, Z.-R. (2019). A technique for delineating the unfolding requirements for substrate entry into retrotranslocons during endoplasmic reticulum-associated degradation. J. Biol. Chem. 294, 20084–20096.

Stein, A., Ruggiano, A., Carvalho, P., and Rapoport, T.A. (2014). Key steps in ERAD of luminal ER proteins reconstituted with purified components. Cell 158, 1375–1388.

Stolz, A., Schweizer, R.S., Schäfer, A., and Wolf, D.H. (2010). Dfm1 forms distinct complexes with Cdc48 and the ER ubiquitin ligases and is required for ERAD. Traffic 11, 1363–1369.

Tirosh, B., Furman, M.H., Tortorella, D., and Ploegh, H.L. (2003). Protein unfolding is not a prerequisite for endoplasmic reticulum-to-cytosol dislocation. J. Biol. Chem. 278, 6664–6672.

Tsuchiya, H., Ohtake, F., Arai, N., Kaiho, A., Yasuda, S., Tanaka, K., and Saeki, Y. (2017). In Vivo Ubiquitin Linkage-type Analysis Reveals that the Cdc48-Rad23/Dsk2 Axis Contributes to K48-Linked Chain Specificity of the Proteasome. Mol. Cell 66, 488–502.e7.

Tyanova, S., Temu, T., and Cox, J. (2016). The MaxQuant computational platform for mass spectrometry-based shotgun proteomics. Nat. Protoc. 11, 2301–2319.

Vasic, V., Denkert, N., Schmidt, C.C., Riedel, D., Stein, A., and Meinecke, M. (2020). Hrd1 forms the retrotranslocation pore regulated by auto-ubiquitination and binding of misfolded proteins. Nat. Cell Biol. 22, 274–281.

Wang, Y., Zhang, Y., and Ha, Y. (2006). Crystal structure of a rhomboid family intramembrane protease. Nature 444, 179–180.

Wolf, D.H., and Stolz, A. (2012). The Cdc48 machine in endoplasmic reticulum associated protein degradation. Biochim. Biophys. Acta 1823, 117–124.

Wu, X., and Rapoport, T.A. (2018). Mechanistic insights into ER-associated protein degradation. Curr. Opin. Cell Biol. 53, 22–28.

Wu, X., Siggel, M., Ovchinnikov, S., Mi, W., Svetlov, V., Nudler, E., Liao, M., Hummer, G., and Rapoport, T.A. (2020). Structural basis of ER-associated protein degradation mediated by the Hrd1 ubiquitin ligase complex. Science 368.

Wu, Z., Yan, N., Feng, L., Oberstein, A., Yan, H., Baker, R.P., Gu, L., Jeffrey, P.D., Urban, S., and Shi, Y. (2006). Structural analysis of a rhomboid family intramembrane protease reveals a gating mechanism for substrate entry. Nat. Struct. Mol. Biol. 13, 1084–1091.

Yasuda, S., Tsuchiya, H., Kaiho, A., Guo, Q., Ikeuchi, K., Endo, A., Arai, N., Ohtake, F., Murata, S., Inada, T., et al. (2020). Stress- and ubiquitylation-dependent phase separation of the proteasome. Nature 578, 296–300.

Ye, Y., Meyer, H.H., and Rapoport, T.A. (2001). The AAA ATPase Cdc48/p97 and its partners transport proteins from the ER into the cytosol. Nature 414, 652–656.

Ye, Y., Shibata, Y., Yun, C., Ron, D., and Rapoport, T.A. (2004). A membrane protein complex mediates retro-translocation from the ER lumen into the cytosol. Nature 429, 841–847.

