## Supplementary material for "The derlin Dfm1 promotes retrotranslocation of folded protein domains from the endoplasmic reticulum": Figure S1

### Fonseca & Carvalho- Figure S1

A)

| | | | | | |
|---|---|---|---|---|---|
| 10 | 20 | 30 | 40 | 50 | 60 |
| MFFNRLSAGK | LLVPLSVVLY | ALFVVILPLQ | NSFHSSNVLV | RGADDVERPA | SYPYDVPDYA |
| 70 | 80 | 90 | 100 | 110 | 120 |
| GYPYDVPDYA | GSYPYDVPDY | AGSSRMNGTE | GPNFYGPFSN | KTVDGSMISL | IAALAVDRVI |
| 130 | 140 | 150 | 160 | 170 | 180 |
| GMENAMPWNL | PADLAWFKRN | TLNKPVIMGR | HTWESIGRPL | PGRKNIILSS | QPGTDDRVTW |
| 190 | 200 | 210 | 220 | 230 | 240 |
| VKSVDEAIAA | CGDVPEIMVI | GGGRVYEQFL | PKAQKLYLTH | IDAEVEGDTH | FPDYEPDDWE |
| 250 | 260 | 270 | 280 | 290 | 300 |
| SVFSEFHDAD | AQNSHSYCFE | ILERRGPSGS | GQVLIKYDYP | KSSFFKKPLS | IACYIFTALM |
| 310 | 320 | 330 | 340 | 350 | 360 |
| GVFVLKTLNM | NVTNRTLQVD | GAGAGAGAGA | GTPKDPAKPP | ATAQVVGWPP | VRSYRKNVMV |
| 370 | 380 | 390 | 400 | 410 | 420 |
| SCQKSSGGPE | AAAFVKVSPG | PKDPAKPPAT | AQVVGWPPVR | SYRKNVMVSC | QKSSGGPEAA |
| 430 | 440 | 450 | 460 | 470 | 480 |
| AFVKVSPGPK | DPAKPPATAQ | VVGWPPVRSY | RKNVMVSCQK | SSGGPEAAAF | VKVSPGPKDP |
| 490 | 500 | 510 | 520 | 530 | 540 |
| AKPPATAQVV | GWPPVRSYRK | NVMVSCQKSS | GGPEAAAFVK | VSPGASGEQK | LISEEDLNGE |
| 550 | 560 | 570 | 580 | 590 | 600 |
| QKLISEEDLN | GEQKLISEED | LNGSSRGEQK | LISEEDLNGE | QKLISEEDLN | GEQKLISEED |
| 610 | 620 | 630 | 640 | | |
| LNGSSRGEQK | ISEEDLNGE | QKLISEEDLN | GEQKLISEED | LNGSTS | |

L-fragment
S-fragment

B)

| Sequence | Start position | End position | Peak Intensity WT | Peak Intensity *Δdfm1* |
|---|---|---|---|---|
| RNTLNKPVIMGR | 139 | 150 | 1.58E+07 | 2.05E+07 |
| NTLNKPVIMGR | 140 | 150 | 2.92E+08 | 1.85E+09 |
| HTWESIGRPLPGR | 151 | 163 | 3.21E+07 | 4.71E+08 |
| HTWESIGRPLPGRK | 151 | 164 | 1.32E+08 | 6.25E+08 |
| KNIILSSQPGTDDR | 164 | 177 | 5.00E+07 | 2.32E+07 |
| KNIILSSQPGTDDRVTWVK | 164 | 182 | 5.10E+07 | 2.56E+08 |
| NIILSSQPGTDDR | 165 | 177 | 5.56E+08 | 1.42E+09 |
| NIILSSQPGTDDRVTWVK | 165 | 182 | 1.91E+09 | 3.81E+09 |
| SVDEAIAACGDVPEIMVIGGGR | 183 | 204 | 4.86E+08 | 8.93E+07 |
| VYEQFLPK | 205 | 212 | 9.44E+08 | 2.32E+09 |
| RGPSGSGQVLIK | 265 | 276 | 1.07E+09 | 2.75E+09 |
| GPSGSGQVLIK | 266 | 276 | 0 | 6.54E+08 |
| TLNMNVTNR | 307 | 315 | 5.66E+08 | 1.26E+09 |
| TLQVDGAGAGAGAGAGTPK | 316 | 334 | 9.81E+08 | 5.18E+07 |
| TLQVDGAGAGAGAGAGTPKDPAKPPATAQVVGWPPVR | 316 | 352 | 2.13E+07 | 0 |
| DPAKPPATAQVVGWPPVR | 335 | 352 | 5.04E+07 | 0 |
