## Supplementary figures and images for "The derlin Dfm1 promotes retrotranslocation of folded protein domains from the endoplasmic reticulum"

### Figure S2

#### Fonseca & Carvalho- Figure S2

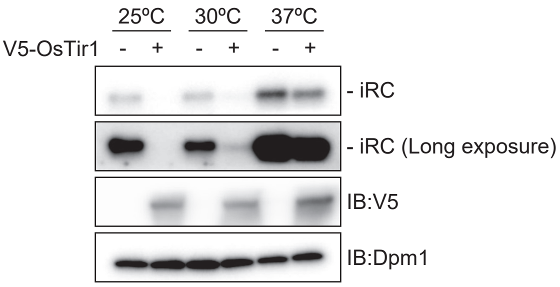

### Figure S3

Fonseca & Carvalho- Figure S3

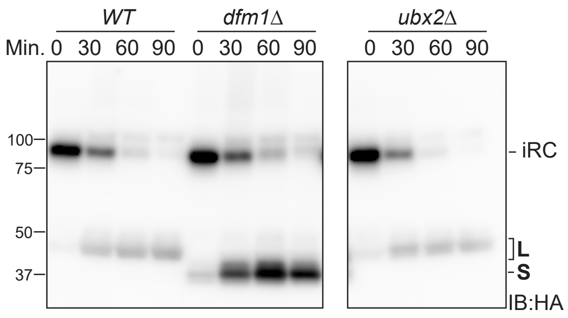

### Figure S4

Fonseca & Carvalho- Figure S4

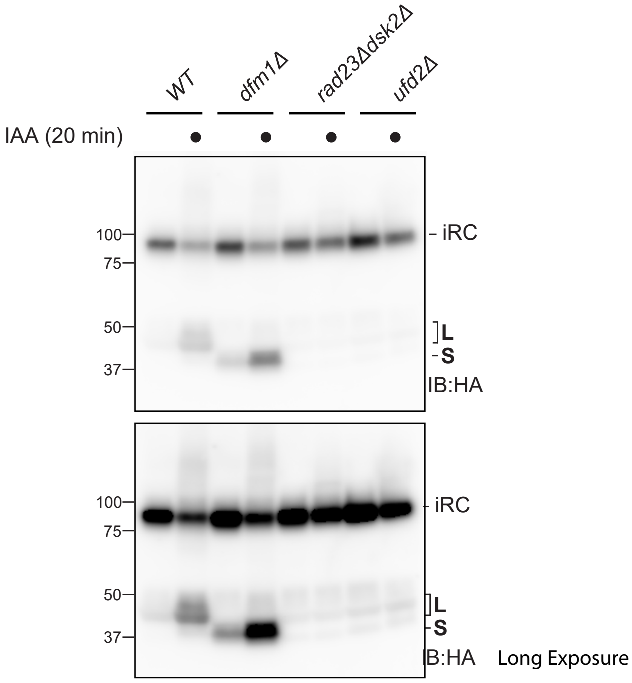
