## Supplemental Tables for "The derlin Dfm1 promotes retrotranslocation of folded protein domains from the endoplasmic reticulum"

**Table S1- List of yeast strains used in this study**

| Strain | Genotype | Reference |
|---|---|---|
| yPC1507 | Mata FY251 ura3-52 his3D200 leu2D1 trp1D62 | This study |
| yPC11111 | Mata URA::GPDp-atTir1-9myc HO::CPYp-3xHA-DHFR-OST(TM)-4aid-9myc::HYGR Asi1::NAT ura3-52 his3D200 leu2D1 trp1D63 | This study |
| yPC11120 | Mata URA::GPDp-atTir1-9myc HO::CPYp-3xHA-DHFR-OST(TM)-4aid-9myc::HYGR Doa10::HIS ura3-52 his3D200 leu2D1 trp1D64 | This study |
| yPC11124 | Mata URA::GPDp-atTir1-9myc HO::CPYp-3xHA-DHFR-OST(TM)-4aid-9myc::HYGR HRD1::CRISPR ura3-52 his3D200 leu2D1 trp1D65 | This study |
| yPC11129 | Mata URA::GPDp-atTir1-9myc Dfm1::KAN HO::CPYp-3xHA-DHFR-OST(TM)-4aid-9myc::HYGR ura3-52 his3D200 leu2D1 trp1D66 pRS415 | This study |
| yPC11130 | Mata URA::GPDp-atTir1-9myc Dfm1::KAN HO::CPYp-3xHA-DHFR-OST(TM)-4aid-9myc::HYGR ura3-52 his3D200 leu2D1 trp1D67 <bPC521> | This study |
| yPC11131 | Mata URA::GPDp-atTir1-9myc Dfm1::KAN HO::CPYp-3xHA-DHFR-OST(TM)-4aid-9myc::HYGR ura3-52 his3D200 leu2D1 trp1D68 <bPC522> | This study |
| yPC11132 | Mata URA::GPDp-atTir1-9myc Dfm1::KAN HO::CPYp-3xHA-DHFR-OST(TM)-4aid-9myc::HYGR ura3-52 his3D200 leu2D1 trp1D69 <bPC1790> | This study |
| yPC11143 | Mata URA::GPDp-atTir1-9myc HO::CPYp-3xHA-DHFR-OST(TM)-4aid-9myc::HYGR ubc7::KAN ura3-52 his3D200 leu2D1 trp1D70 | This study |
| yPC11167 | Mata URA::GPDp-atTir1-9myc HO::CPYp-3xHA-DHFR( CYS1&CYS2)-OST(TM)-4aid-9myc::HYGR ura3-52 his3D200 leu2D1 trp1D71 | This study |
| yPC11168 | Mata URA::GPDp-atTir1-9myc Dfm1::KAN HO::CPYp-3xHA-DHFR( CYS1&CYS2)-OST(TM)-4aid-9myc::HYGR ura3-52 his3D200 leu2D1 trp1D72 | This study |
| yPC11191 | Mata URA::GPDp-atTir1-9myc HO::CPYp-3xHA-DHFR-OST(TM)-4aid-9myc::HYGR ura3-52 his3D200 leu2D1 trp1D73 pRS415 | This study |
| yPC11192 | Mata URA::GPDp-atTir1-9myc HO::CPYp-3xHA-DHFR-OST(TM)-4aid-9myc::HYGR ura3-52 his3D200 leu2D1 trp1D74 <bPC521> | This study |
| yPC11193 | Mata URA::GPDp-atTir1-9myc HO::CPYp-3xHA-DHFR-OST(TM)-4aid-9myc::HYGR ura3-52 his3D200 leu2D1 trp1D75 <bPC522> | This study |
| yPC11194 | Mata URA::GPDp-atTir1-9myc HO::CPYp-3xHA-DHFR-OST(TM)-4aid-9myc::HYGR ura3-52 his3D200 leu2D1 trp1D76 <bPC1790> | This study |
| yPC11203 | Mata URA::GPDp-atTir1-9myc HO::CPYp-3xHA-DHFR( CYS1)-OST(TM)-4aid-9myc::HYGR ura3-52 his3D200 leu2D1 trp1D77 | This study |
| yPC11204 | Mata URA::GPDp-atTir1-9myc Dfm1::KAN HO::CPYp-3xHA-DHFR( CYS1)-OST(TM)-4aid-9myc::HYGR ura3-52 his3D200 leu2D1 trp1D78 | This study |

| | | |
|---|---|---|
| yPC11205 | Mata URA::GPDp-atTir1-9myc HO::CPYp-3xHA-DHFR( CYS2)-OST(TM)-4aid-9myc::HYGR ura3-52 his3D200 leu2D1 trp1D79 | This study |
| yPC11206 | Mata URA::GPDp-atTir1-9myc Dfm1::KAN HO::CPYp-3xHA-DHFR( CYS2)-OST(TM)-4aid-9myc::HYGR ura3-52 his3D200 leu2D1 trp1D80 | This study |
| yPC11233 | Mata URA::GPDp-atTir1-9myc HO::CPYp-3xHA-DHFR-OST(TM)-4aid-9myc::HYGR ura3-52 his3D200 leu2D1 trp1D81 | This study |
| yPC11234 | Mat? Asi1::NAT Doa10::HIS Hrd1::KAN URA::GPDp-atTir1-9myc HO::CPYp-3xHA-DHFR-OST(TM)-4aid-9myc::HYGR ura3-52 his3D200 leu2D1 trp1D82 | This study |
| yPC11236 | Mata URA::GPDp-atTir1-9myc Der1::HIS HO::CPYp-3xHA-DHFR-OST(TM)-4aid-9myc::HYGR ura3-52 his3D200 leu2D1 trp1D83 | This study |
| yPC11237 | Mata URA::GPDp-atTir1-9myc Dfm1::KAN HO::CPYp-3xHA-DHFR-OST(TM)-4aid-9myc::HYGR ura3-52 his3D200 leu2D1 trp1D84 | This study |
| yPC11238 | Mata URA::GPDp-atTir1-9myc Der1::HIS Dfm1::KAN HO::CPYp-3xHA-DHFR-OST(TM)-4aid-9myc::HYGR ura3-52 his3D200 leu2D1 trp1D85 | This study |
| yPC11242 | Mata URA::GPDp-atTir1-9myc 3xHA-DHFR(29-31_Pro)-OST(TM)-4aid-9myc::HYGR ura3-52 his3D200 leu2D1 trp1D86 | This study |
| yPC11246 | Mata URA::GPDp-atTir1-9myc Dfm1::KAN 3xHA-DHFR(29-31_Pro)-OST(TM)-4aid-9myc::HYGR ura3-52 his3D200 leu2D1 trp1D87 | This study |
| yPC11271 | Mata URA::GPDp-atTir1-9myc HO::CPYp-3xHA-DHFR-OST(TM)-4aid-9myc::HYGR DFM(1-283-SBP)::CRISPR ura3-52 his3D200 leu2D1 trp1D92 | This study |
| yPC11280 | Mat? Asi1::NAT DOA10::HIS HRD1::KAN URA::GPDp-atTir1-9myc HO::CPYp-3xHA-DHFR-OST(TM)-4aid-9myc::HYGR DFM1::CRISPR ura3-52 his3D200 leu2D1 trp1D93 | This study |
| yPC11282 | Mat? Asi1::NAT DOA10::HIS HRD1::KAN URA::GPDp-atTir1-9myc HO::CPYp-3xHA-DHFR-OST(TM)-4aid-9myc::HYGR Der1::CRISPR ura3-52 his3D200 leu2D1 trp1D94 | This study |
| yPC11287 | Mata URA::GPDp-osTir1-V5 HO::CPYp-3xHA-DHFR-OST(TM)-4aid-9myc::HYGR ura3-52 his3D200 leu2D1 trp1D95 | This study |
| yPC11288 | Mata URA::GPDp-osTir1-V5 HO::CPYp-3xHA-DHFR-OST(TM)-4aid-9myc::HYGR DFM1:KAN ura3-52 his3D200 leu2D1 trp1D96 | This study |
| yPC11289 | Mat? URA::GPDp-osTir1-V5 HO::CPYp-3xHA-DHFR-OST(TM)-4aid-9myc::HYGR npl4-1 ura3-52 his3D200 leu2D1 trp1D97 | This study |
| yPC11678 | Mat? URA::GPDp-osTir1-V5 HO::CPYp-3xHA-DHFR-OST(TM)-4aid-9myc::HYGR DFM1::KAN npl4-1 ura3-52 his3D200 leu2D1 trp1D98 | This study |
| yPC11147 | Mat? URA::GPDp-osTir1-V5 HO::CPYp-3xHA-DHFR-OST(TM)-4aid-9myc::HYGR cdc48-6 ura3-52 his3D200 leu2D1 trp1D99 | This study |
| yPC11148 | Mat? URA::GPDp-osTir1-V5 HO::CPYp-3xHA-DHFR-OST(TM)-4aid-9myc::HYGR DFM1::KAN cdc48-6 ura3-52 his3D200 leu2D1 trp1D100 | This study |

| | | |
|---|---|---|
| yPC11681 | MATa HO::ADHp-3xHA-DHFR-OST(TM)-4aid-9myc::HYGR URA::GPDp-atTir1-9myc ura3-52 his3D200 leu2D1 trp1D101 | This study |
| yPC11682 | MATa HO::ADHp-3xHA-DHFR-OST(TM)-4aid-9myc::HYGR URA::GPDp-atTir1-9myc DFM1:KAN ura3-52 his3D200 leu2D1 trp1D102 | This study |
| yPC11151 | MAT? HO::ADHp-3xHA-DHFR-OST(TM)-4aid-9myc::HYGR URA::GPDp-atTir1-9myc Rad23::HIS Dsk2::KAN ura3-52 his3D200 leu2D1 trp1D103 | This study |
| yPC11155 | MATa HO::ADHp-3xHA-DHFR-OST(TM)-4aid-9myc::HYGR URA::GPDp-atTir1-9myc Ubx2::KAN ura3-52 his3D200 leu2D1 trp1D104 | This study |
| yPC11156 | MATa HO::ADHp-3xHA-DHFR-OST(TM)-4aid-9myc::HYGR URA::GPDp-atTir1-9myc ufd2::KAN ura3-52 his3D200 leu2D1 trp1D105 | This study |

**Table S2- List of plasmids used in this study**

| Plasmid | Insert | Reference |
|---|---|---|
| PB565 | pRS425-Gal1p | This study |
| bPC1614 | pMK27-GPD_atTir1-9xMyc | M Morawska et al 2013 |
| bPC1680 | pFA6a_CPYp-3xHA-DHFR-OST1(TM)-4xAID-9xMyc_HphMX6 | This study |
| bPC1670 | pFA6a_3xHA-DHFR(29-31PRO)-OST1(TM)-4xAID-9xMyc_HphMX6 | This study |
| bPC521 | pRS425-Gal1p_Dfm1-Flag | This study |
| bPC522 | pRS425-Gal1p_Dfm1(WR-AA)-Flag | This study |
| bPC1790 | pRS425-Gal1p_Dfm1(GxxxG-AA)-Flag | This study |
| bPC1791 | pRS306-GPDp_osTir1-V5 | This study |
| bPC1828 | pFA6a_CPYp_3xHA-DHFR-CYS1-OST1(TM)-4xAID-9xMyc_HphMX6 | This study |
| bPC1829 | pFA6a_CPYp_3xHA-DHFR-CYS2-OST1(TM)-4xAID-9xMyc_HphMX6 | This study |
| bPC1830 | pFA6a_CPYp_3xHA-DHFR-CYS1&2-OST1(TM)-4xAID-9xMyc_HphMX6 | This study |
| bPC1681 | pML107_GAPp-Cas9_Dfm1-gRNA1 | This study |
| bPC1695 | pML107_GAPp-Cas9_Hrd1-gRNA1 | This study |
| bPC1827 | pFA6a_ADHp_3xHA-DHFR-OST1(TM)-4xAID-9xMyc_HphMX6 | This study |
| bPC1798 | pFA6a_ADHp_3xHA-yeGFP-DHFR-OST1(TM)-4xAID-9xMyc_HphMX6 | This study |
